# Dysregulated nuclear Lamin B1 in DYT1 dystonia thickens the nuclear lamina and disrupts 14-3-3 proteins

**DOI:** 10.1101/2025.07.09.662391

**Authors:** Yuntian Duan, Masood Sepehrimanesh, Md Abir Hosain, Haochen Cui, Jacob Stagray, Xinggui Shen, Ying Xiao, Yuqing Li, Chun-Li Zhang, Baojin Ding

## Abstract

Childhood-onset DYT1 dystonia is caused by a heterozygous ΔE mutation in the *TOR1A* gene, which encodes a membrane-embedded AAA+ (ATPase Associated with diverse cellular Activities) ATPase. However, the mechanism by which ΔE induces dystonia remains poorly understood. Previously, using patient-derived neurons, we identified dysregulation of nuclear Lamin B1, at both expression levels and subcellular distribution, as a key contributor to DYT1 pathology. In the present study, we utilized DYT1 patient fibroblast cells and induced human neurons to investigate the molecular basis and consequences of Lamin B1 dysregulation. We found that elevated nuclear Lamin B1 thickens the nuclear lamina and deforms the nucleus, impairing nucleocytoplasmic transport. Proteomic analysis of human iPSC-derived neurons revealed that mislocalized Lamin B1 disrupts essential signaling pathways involved in neuronal function. Notably, 14-3-3 proteins, abundant brain molecular chaperones critical for neuronal development and homeostasis, were the most strongly associated with mislocalized Lamin B1. Functional studies showed that downregulation of 14-3-3 proteins impairs neurodevelopment in healthy neurons, while their upregulation rescues DYT1 neuronal defects by reducing Lamin B1 mislocalization. These findings elucidate a mechanistic link between nuclear deformation and cellular dysfunction in DYT1 dystonia and highlight Lamin B1 and 14-3-3 proteins as potential therapeutic targets.

## INTRODUCTION

The nuclear lamina is a filamentous meshwork lining inner nuclear membrane (INM) in eukaryotic cells, primarily composed of A-type (lamin A, lamin C, and minor isoforms) and B-type (lamin B1 and lamin B2) lamins ^1,2^. It provides structural support, regulates chromatin organization, and is essential for nuclear stability, gene expression, and cell signaling ^3–8^. Dysregulation of nuclear lamina has been implicated in a range of conditions, including muscular dystrophy, cardiomyopathy, aging, cancer, and neurological diseases ^2,9,10^. Such dysregulation may arise from mutations in lamin-encoding genes, altered expression levels, or protein mislocalization ^11–13^. Nuclear Lamin B1, a major components of B-type Lamins, is ubiquitously expressed in human cells ^14,15^. Mutations in the *LMNB1* gene or abnormal expression of Lamin B1 can significantly disrupt cellular function and are linked to aging and various diseases, including neurological and muscular disorders ^16–21^. However, the mechanisms by which nuclear lamina dysregulation contributes to cellular dysfunction, particularly in the nervous system, remain poorly understood.

DYT1 dystonia is a neurodevelopmental movement disorder that typically manifests in childhood and adolescence, a critical period for motor learning ^22–24^. Most cases are caused by a heterozygous 3-bp deletion (ΔGAG) in the *TOR1A* gene, which eliminates a glutamate residue in the C-terminal region of the TorsinA protein (ΔE) ^25–28^. Torsins are members of the evolutionarily conserved AAA+ ATPase superfamily, which use ATP hydrolysis to disassemble protein complexes in crucial biological processes, including membrane trafficking, cytoskeleton dynamics, vesicle fusion and stress to responses ^29–32^. TorsinA is primarily localized to the lumen of the endoplasmic reticulum (ER) and nuclear envelope (NE) ^33–35^. In the ER, it contributes to protein processing, the ER stress response, and ER-associated protein degradation ^36–41^. At the NE, TorsinA interacts with the KASH domain of nesprins to link the NE to the cytoskeleton ^42^. It also regulates NE morphology ^35,43–45^, nuclear pore complex localization and maturation ^44–48^, and the nuclear egress of large ribonucleoprotein granules ^49^ and herpes simplex virus ^50^.

The ΔE mutation results in TorsinA loss-of-function ^51–54^, but how this leads to dystonia remains unclear. Studies in mice ^22,48,53,55^, *C. elegans*^47^, *Drosophila* ^49^, and mammalian cells ^35,43,44,46,56^ have demonstrated that the loss of torsin causes abnormal NE morphology and cellular dysfunction. However, the molecular mechanisms linking TorsinA dysfunction to NE abnormalities and their role in DYT1 pathophysiology remain to be fully elucidated.

Previously, we modeled DYT1 dystonia using patient-derived neurons generated by direct conversion of fibroblasts or by induction and differentiation of patient-derived induced pluripotent stem cells (iPSCs) ^44,57–61^. These patient-specific neurons, carrying the heterozygous *TOR1A* ΔE mutation, exhibited disease-relevant cellular deficits, including abnormal NE morphology, impaired neurodevelopment, and disrupted nucleocytoplasmic transport (NCT) ^44,62^. A striking finding was the dysregulation of nuclear Lamin B1 at both expression levels and subcellular distribution in human DYT1 neurons ^44,63^. This dysregulation could be reproduced by ectopic expression of the ΔE mutant or by shRNA-mediated knockdown of endogenous *TOR1A* in healthy neurons. Notably, reducing Lamin B1 levels significantly rescued the cellular phenotypes in DYT1 neurons, suggesting that Lamin B1 dysregulation may represent a key molecular mechanism driving DYT1 pathology ^44^.

To better understand how dysregulated Lamin B1 contributes to cellular deficits in DYT1 cells, we hypothesized that elevated Lamin B1 levels alter nuclear lamina structure, while mislocalized cytoplasmic Lamin B1 sequesters key factors and signaling pathways critical for neuronal function. In this study, ultrastructural analysis of the nuclear envelope in DYT1 cells revealed that increased nuclear Lamin B1 thickens the nuclear lamina and deforms the nucleus, leading to impaired nucleocytoplasmic transport. Proteomic analyses further showed that mislocalized Lamin B1 disrupts numerous factors and pathways essential for various biological processes and neuron functions. Among these, 14-3-3 proteins, molecular chaperones highly expressed in the brain and critical for neuronal function ^64,65^, were strongly associated with mislocalized Lamin B1. Importantly, overexpression of 14-3-3 proteins in DYT1 neurons significantly reduced Lamin B1 mislocalization and rescued neuronal deficits. These findings address a long-standing question regarding nuclear deformation in DYT1 cells and provide new insight into the pathogenic role of Lamin B1 dysregulation, highlighting potential therapeutic targets for DYT1 dystonia.

## RESULTS

### Nuclear LMNB1 dysregulation in DYT1 patient fibroblasts

Previously, we demonstrated dysregulation of nuclear Lamin B1 in both expression and subcellular localization in DYT1 patient-derived neurons ^44^. To test whether this dysregulation also occurs in patient fibroblast cells, we obtained four DYT1 patient fibroblast cell lines along with sex- and age-matched healthy controls. All DYT1 patient cell lines were chosen according to stringent criteria: (i) the donors were clinically diagnosed with childhood-onset dystonia, and (ii) their condition attributed to the heterozygous GAG deletion in the *TOR1A* gene. The presence of the heterozygous GAG deletion in exon 5 of the *TOR1A* gene was confirmed through DNA sequencing (**Fig. 1A**). Firstly, we examined nuclear Lamins via immunocytochemistry (ICC). Healthy fibroblasts displayed regular nuclear morphology and even distribution of both nuclear Lamin A/C and Lamin B1. In contrast, DYT1 fibroblasts frequently exhibited wrinkles, infoldings and bright foci (**Fig. 1B**). While the cytoplasmic nuclear Lamin B1 in DYT1 fibroblasts tend to be higher than in healthy controls, the mislocalized Lamin B1 was not as obvious as in patient-derived neurons ^44^. This difference is likely due to the diluted mislocalized nuclear Lamin B1 in the relatively larger size of cytoplasm in fibroblasts. In DYT1 neurons, the mislocalized Lamin B1 can be accumulated in neuron processes during differentiation and maturation, resulting in more pronounced Lamin B1 mislocalizaiton ^44,63^.

**Figure 1.**
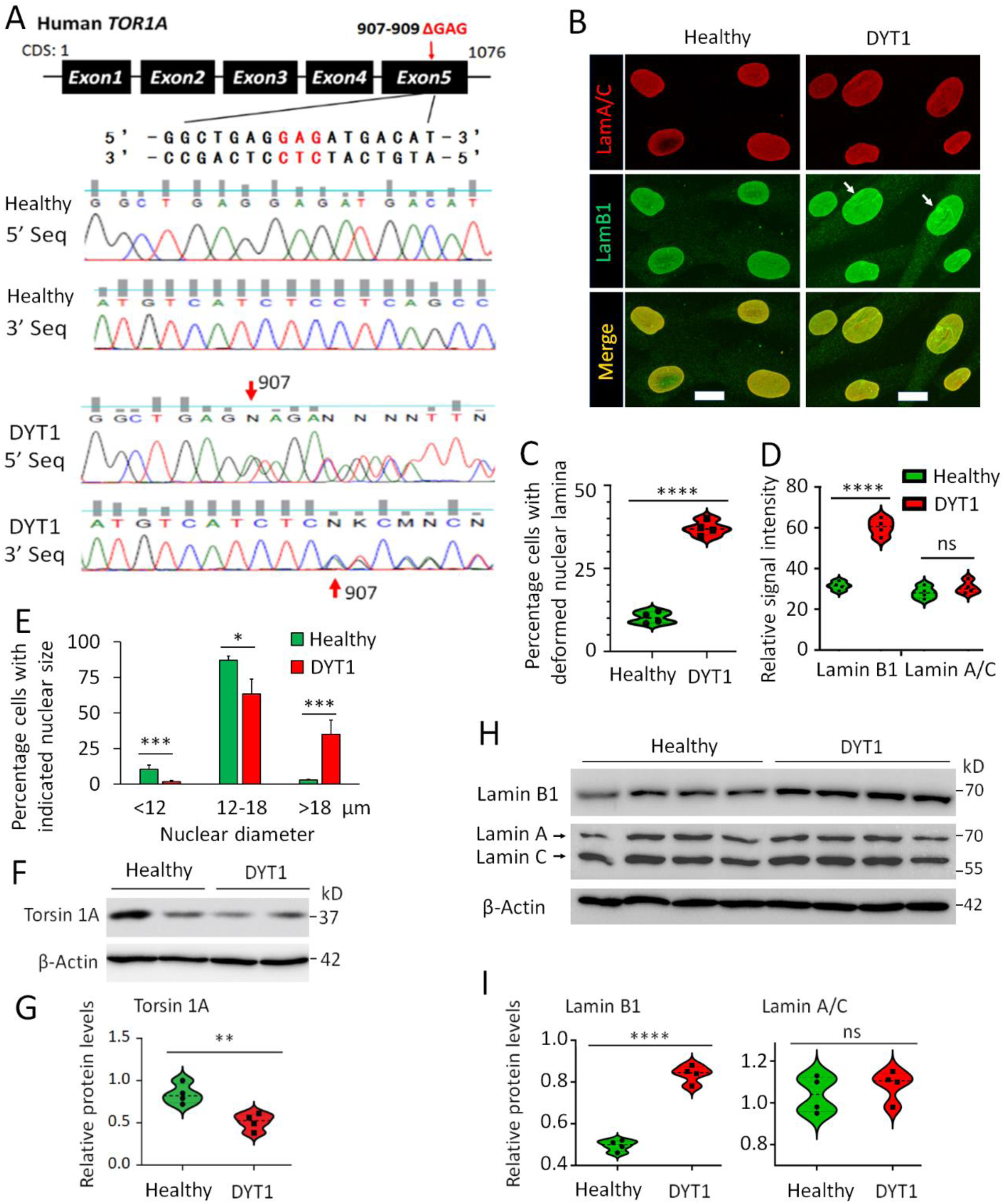
Nuclear Lamin B1 is dysregulated in DYT1 patient fibroblast cells. (**A**) Heterozygous GAG deletion in the *TOR1A* gene in DYT1 patient-derived fibroblasts. (Top) Schematic of the human *TOR1A* gene highlighting the GAG deletion site and flanking DNA sequence. (Bottom) Forward (5′) and reverse (3′) sequencing results of PCR amplicons from healthy control (GM00024) and DYT1 fibroblast cells (GM03211). DYT1 cells show one wild-type allele and one allele with a shifted sequence starting at the GAG deletion site (indicated by red arrows). (**B**) Representative confocal micrographs of immunocytochemistry (ICC) showing the nuclear lamina in fibroblasts from a healthy control (GM00024) and a DYT1 patient (GM03211). Arrows indicate disrupted nuclear lamina and abnormal nuclear envelope morphology. Scale bars: 10 µm. (**C**) Quantification of the percentage of cells with deformed nuclear lamina based on ICC analysis in panel (B). Each dot represents the mean value for one cell line, with >100 cells analyzed per line. (**D**) Quantification of relative signal intensity of Lamin B1 and Lamin A/C from ICC results in panel (B). Each dot represents the mean value for one cell line, with >100 cells analyzed per line. (**E**) Distribution of nuclear size (diameter) in healthy and DYT1 patient fibroblasts. *N* (subjects) = 4 per group. *n* (nuclei) = 246 for healthy controls and 231 for DYT1 cells, from three independent experiments. (**F**) Western blot showing TorsinA protein levels in healthy and DYT1 fibroblasts. (**G**) Quantification of TorsinA protein levels in panel (F), normalized to actin. (**H**) Western blots showing protein levels of nuclear Lamin B1 and Lamin A/C in healthy and DYT1 fibroblasts. (**I**) Quantification of Lamin B1 and Lamin A/C protein levels in panel (H), normalized to actin. For C-E, G, and I, ns, no significant difference; * p < 0.05; ** p < 0.01; *** p < 0.001; **** p < 0.0001. Student’s t-test.

We then conducted a careful examination of nuclear morphology and found that a significantly higher number of DYT1 cells exhibited deformed nucleus, defined with wrinkles and infoldings (**Fig. 1C**). Quantification of signal intensity indicated that the expression level of nuclear Lamin B1 in DYT1 cells is significantly higher than that in healthy controls, while no significant difference was observed for nuclear Lamin A/C (**Fig. 1D**). Moreover, DYT1 fibroblasts tend to have larger nuclei, with approximately 35% of cells exhibiting nuclear diameters greater than 18 µm. In contrast, this fraction was less than 3% in healthy controls, in which the nuclear size of the majority (>95%) cells is between 12 to 18 µm (**Fig. 1E**). The abnormal nuclear size observed in DYT1 cells may be related to disrupted nuclear transport ^66,67^. Furthermore, we analyzed protein expression levels. TorsinA protein was found to be significantly lower in DYT1 patient fibroblast cells compared with healthy controls (**Fig. 1F and G**). Consistent with ICC data showing a stronger nuclear Lamin B1 signal in DYT1 cells (**Fig. 1B and D**), the protein level of nuclear Lamin B1 was notably higher than in healthy controls, while no significant changes were observed for nuclear Lamin A/C (**Fig. 1H and I**). Taken together, these results indicate that deformed nucleus and dysregulated nuclear lamin B1 also occur in DYT1 patient fibroblasts.

### Lamin B1 dysregulation causes nuclear lamina thickening and deformation

To further investigate the nuclear morphology of DYT1 fibroblasts, we employed transmission electron microscopy (TEM). Consistent with the results obtained from ICC (**Fig. 1B**), DYT1 nuclei displayed rough and rigid outlines in contrast to the smooth and regular nuclear morphology observed in healthy cells (**Fig. 2A and B**). Notably, DYT1 cells exhibited a thickened electron dense area immediately beneath the INM (**Fig. 2B1**). This thickened electron dense area was consistently observed in all four DYT1 patient cell lines (**Fig. 2C and D**), indicating that deformed nuclei and thickened electron dense beneath the INM are common features in DYT1 cells, consistent with previous findings in DYT1 mouse models and reprogrammed DYT1 neurons ^44,53^.

**Figure 2.**
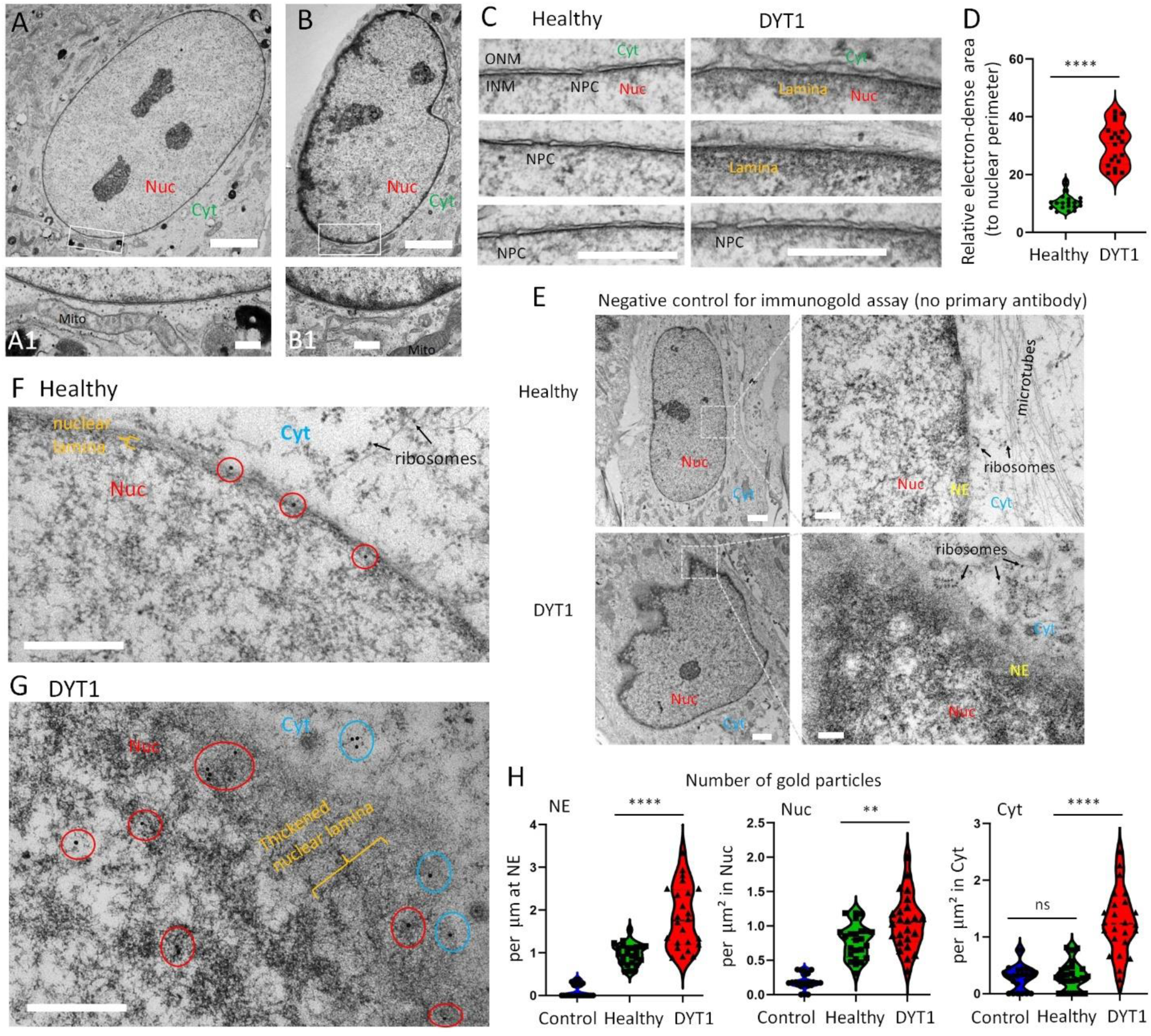
Dysregulated nuclear Lamin B1 causes thickening of the nuclear lamina in DYT1 fibroblast cells. **(A)** Representative transmission electron micrograph (TEM) of a healthy fibroblast cell (GM00024). A higher magnification view of the boxed region is shown in (A1) below. Nuc, nucleus; Cyt, cytoplasm; Mito, mitochondria. Scale bars: 5 µm (A), 500 nm (A1). **(B)** Representative TEM of a DYT1 fibroblast cell (GM03211). A higher magnification view of the boxed region is shown in (B1) below. Scale bars: 5 µm (B), 500 nm (B1). **(C)** High-magnification TEM images focusing on the nuclear envelope in healthy controls (GM 03652, GM04506, AG07473) and DYT1 fibroblast cells (NDS00305, GM02304, NDS00301). INM, inner nuclear membrane; ONM, outer nuclear membrane; NPC, nuclear pore complex. Scale bars: 500 nm. **(D)** Quantification of the electron-dense area immediately beneath the INM, normalized to nuclear perimeter. *N* (nuclei) = 20 per group from three independent experiments. ****p < 0.0001; Student’s *t*-test. **(E)** TEM images of immunogold labeling in healthy (GM00024) and DYT1 (GM03211) fibroblasts processed without primary antibody (negative control). Scale bars: 2 µm (low magnification) and 200 nm (high magnification). Nuc, nucleus; Cyt, cytoplasm; NE, nuclear envelope. **(F)** Immunogold TEM of Lamin B1 in healthy fibroblasts (GM00024). Gold particles marking Lamin B1 localization are circled. Scale bar: 500 nm. **(G)** Immunogold TEM of Lamin B1 in DYT1 fibroblasts GM03211). Gold particles on the nuclear envelope and nucleoplasm are circled in red; cytoplasmic gold particles are circled in blue. Scale bar: 500 nm. **(H)** Quantification of gold particle distribution at the nuclear envelope (NE), in the nucleoplasm (Nuc), and in the cytoplasm (Cyt). Control group represents samples without primary antibody. *n* (nuclei) = 24 per group from three independent experiments. ns, not significant; **p < 0.01; ****p < 0.0001; One-way ANOVA with Dunnett’s multiple comparison test.

To determine whether the dysregulated nuclear Lamin B1 contributes to the thickened electron dense in DYT1 cells, we conducted immunogold analysis using anti-Lamin B1 antibody combined with a secondary antibody conjugated with 15 nm gold particles. Negligible gold particles were observed in negative control samples treated without primary antibody (**Fig. 2E**), confirming the specificity of the immunogold labeling. Compared with the distribution of gold particles, which were almost exclusively localized on the NE in healthy control cells (**Fig. 2F**), a significantly higher number of gold particles can be found in DYT1 cells (**Fig. 2G**). These gold particles were widely distributed in the cytoplasm and nucleoplasm around the NE, with a particularly higher enrichment in the electron-dense area beneath the INM compared to healthy controls (**Fig. 2H**). These results provide strong evidence that the thickened electron-dense area observed in DYT1 cells is, in fact, the thickened nuclear lamina resulting from dysregulated nuclear Lamin B1.

Additionally, we examined the impact of nuclear Lamin B1 levels on NE morphology by ectopically expressing LMNB1 along with a GFP reporter. Remarkably, nearly all cells overexpressing LMNB1 exhibited abnormal NE morphology, where cells overexpressing GFP alone showed no noticeable changes (**Fig. S1A, B and F**). Interestingly, the downregulation of LMNB1 using shRNAs (co-expressing an mCherry reporter) did not visibly affect NE morphology (**Fig. S1C-E**). Consistently, TEM analysis revealed that LMNB1 overexpression notably disrupted NE morphology, showing a rigid outline and thickened electron-dense layer beneath the INM, while no obvious changes were observed in LMNB1 knockdown cells (**Fig. S1H**). These results suggest that the upregulation of nuclear Lamin B1 significantly disrupts NE morphology, whereas its downregulation does not.

### Impaired NCT of both proteins and mRNAs in DYT1 cells

To assess whether NCT activities are disrupted in DYT1 fibroblasts, we utilized a dual reporter system to measure protein nuclear transport. As described previously ^62,68–70^, this reporter system consists of a GFP reporter fused with a nuclear export signal (GFP-NES) and an RFP reporter fused with a nuclear localization signal (RFP-NLS). In cells with normal NCT activity, GFP and RFP are localized in the cytoplasm and nucleus, respectively. However, in cells with compromised NCT, this subcellular distribution will be disrupted (**Fig. 3A**). An increased ratio of nuclear to cytoplasmic GFP (Nuc/Cyt) indicates impaired protein export, while a higher ratio of cytoplasmic to nuclear RFP (Cyt/Nuc) indicates compromised protein import. In healthy fibroblasts, GFP and RFP were almost exclusively localized in the cytoplasm and nucleus, respectively. In contrast, DYT1 fibroblast cells exhibited substantial amount of GFP inside the nucleus and obvious RFP signals in the cytoplasm (**Fig. 3A**). Consistently, the ratios of GFP (Nuc/Cyt) and RFP (Cyt/Nuc) were significantly higher in DYT1 cells compared to healthy controls (**Fig. 3B and C**), indicating impaired protein nuclear export and import activities in DYT1 fibroblasts.

**Figure 3.**
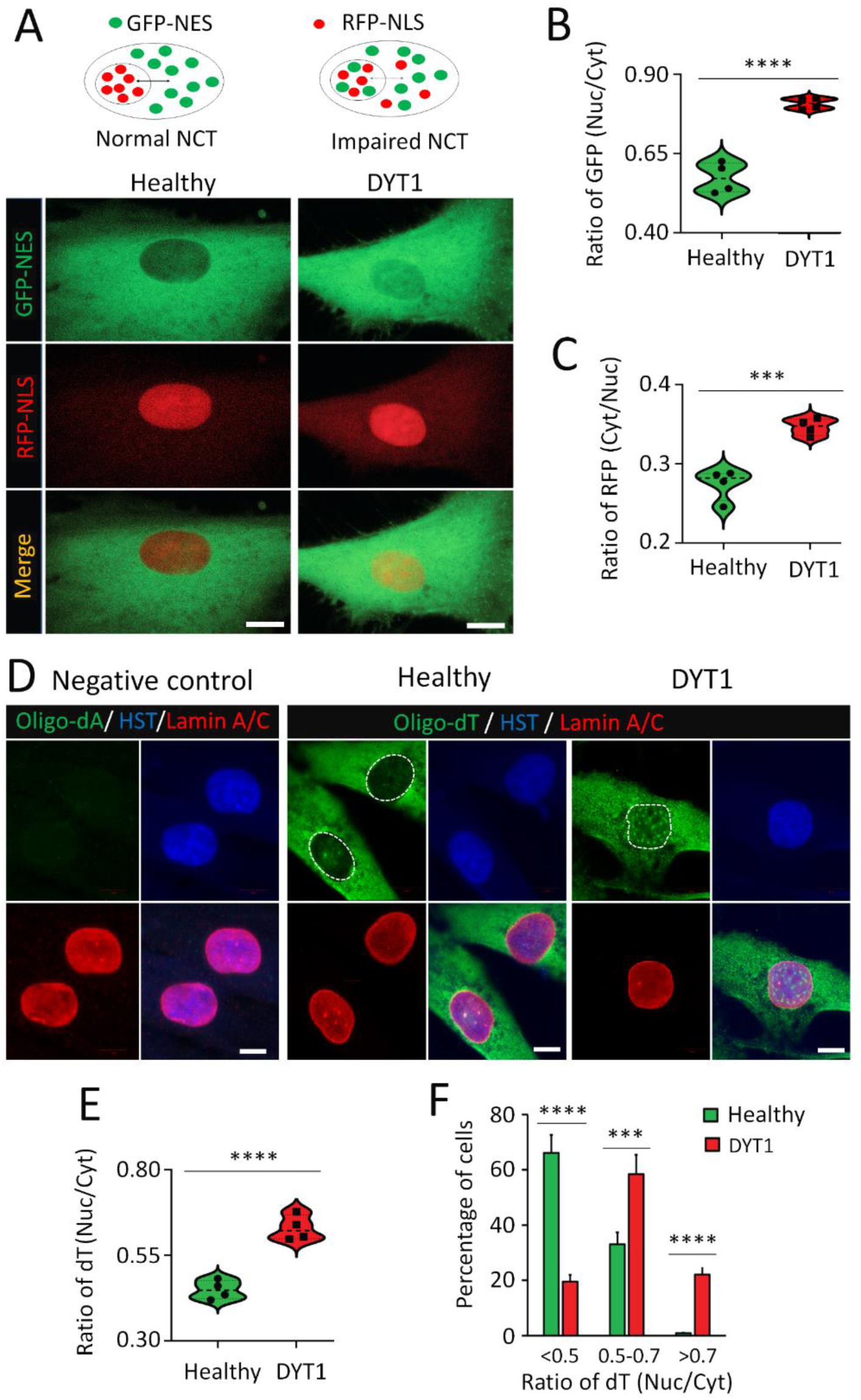
Impaired nucleocytoplasmic transport (NCT) of both protein and mRNA cargos in DYT1 cells. (**A**) Top: Schematic of the dual-reporter system used to assess protein NCT. NES, nuclear export signal; NLS, nuclear localization signal. Bottom: Representative confocal micrographs showing reporter distribution in healthy (GM00024) and DYT1 (GM03211) fibroblasts at 7 days post-infection (dpi). Scale bars: 10 µm. (**B**) Quantification of nuclear-to-cytoplasmic GFP signal ratio for the NES-GFP reporter. *N* (subjects) = 4; *n* (cells) = 173 (healthy) and 162 (DYT1) from three independent experiments. ****p < 0.0001, Student’s *t*-test. (**C**) Quantification of cytoplasmic-to-nuclear RFP signal ratio for the NLS-RFP reporter. *N* (subjects) = 4; *n* (cells) = 173 (healthy) and 162 (DYT1) from three independent experiments. ***p < 0.001, Student’s *t*-test. (**D**) Representative confocal images of fluorescence in situ hybridization (FISH) assay. Oligo-dA probes served as negative control; oligo-dT probes were used to detect endogenous polyadenylated mRNAs. Nuclei were stained with Lamin A/C and Hoechst 33342 (HST). Dotted circles delineate the nuclear boundary. Scale bars: 10 µm. (**E**) Quantification of nuclear-to-cytoplasmic oligo-dT signal ratio, reflecting mRNA distribution. *N* (subjects) = 4; *n* (cells) = 156 (healthy) and 125 (DYT1) from three independent experiments. ****p < 0.0001, Student’s *t*-test. (**F**) Distribution of cells according to the ratio of nuclear to cytoplasmic oligo-dT signal. *N* (subjects) = 4; *n* (cells) = 156 (healthy) and 125 (DYT1) from three independent experiments. ***p < 0.001; ****p < 0.0001; Student’s *t*-test.

We also performed fluorescence *in situ* hybridization (FISH) to examine whether mRNA export was impaired in DYT1 fibroblasts. As previously described ^62,68,70^, digoxigenin (DIG)-labeled oligo-dT probes were used to detect the subcellular distribution of Poly (A) mRNAs. FISH specificity was confirmed by the absence of signals when DIG-labeled oligo-dA was used as a probe (**Fig. 3D**). mRNA nuclear export was quantified by the ratio of nuclear to cytoplasmic oligo-dT signals [dT(Nuc/Cyt)]. In healthy controls, a majority of mRNAs were localized in the cytoplasm. In sharp contrast, a substantial amount of mRNAs retained inside the nucleus of DYT1 cells (**Fig. 3D**). A significantly larger number of DYT1 cells showed a higher ratio of dT(Nuc/Cyt) compared to healthy controls (**Fig. 3E**). For instance, less than <1% of healthy cells exhibited a dT(Nuc/Cyt) ratio greater than 0.7, while in DYT1 cells, this fraction was more than 20% (**Fig. 3F**). Together, dual reporter assay and FISH analysis clearly indicate that both protein NCT and mRNA nuclear export are impaired in DYT1 fibroblasts.

Additionally, we measured the NCT in cells with altered LMNB1 expression (**Fig. S1A-D**). Consistent with the NE morphology results (**Fig. S1F**), only nuclear LMNB1 overexpression significantly disrupted NCT, while LMNB1 knockdown had no notable effect (**Fig. S1G**). These findings indicate that nuclear Lamin B1 levels are critical for maintaining both NE morphology and NCT activity. The NCT impairment observed in DYT1 cells is thus likely linked to nuclear deformation caused by LMNB1 dysregulation.

### Identify proteins disrupted by cytoplasmic Lamin B1 in hiPSC-derived MNs

We have previously demonstrated that nuclear Lamin B1 is progressively accumulated in the cytoplasm and neuron processes in DYT1 patient-derived neurons and it may constitute a prominent pathogenic factor in DYT1 pathogenesis ^44,63^. To investigate the impact of mislocalized Lamin B1 on neuronal function and its potential molecular link to DYT1 dystonia, we propose that the progressive accumulation of cytosolic Lamin B1 may sequester key factors involved in critical signaling pathways, ultimately leading to widespread cellular dysfunction. We performed Co-immunoprecipitation coupled with mass spectrometry (Co-IP/MS) to identify cytosolic Lamin B1-interacting proteins in hiPSC-derived MNs (hiPSC-MNs), as this approach provides high-quality neurons with good yield and purity for biochemical studies (**Fig. 4A**) ^62,71,72^.

**Figure 4.**
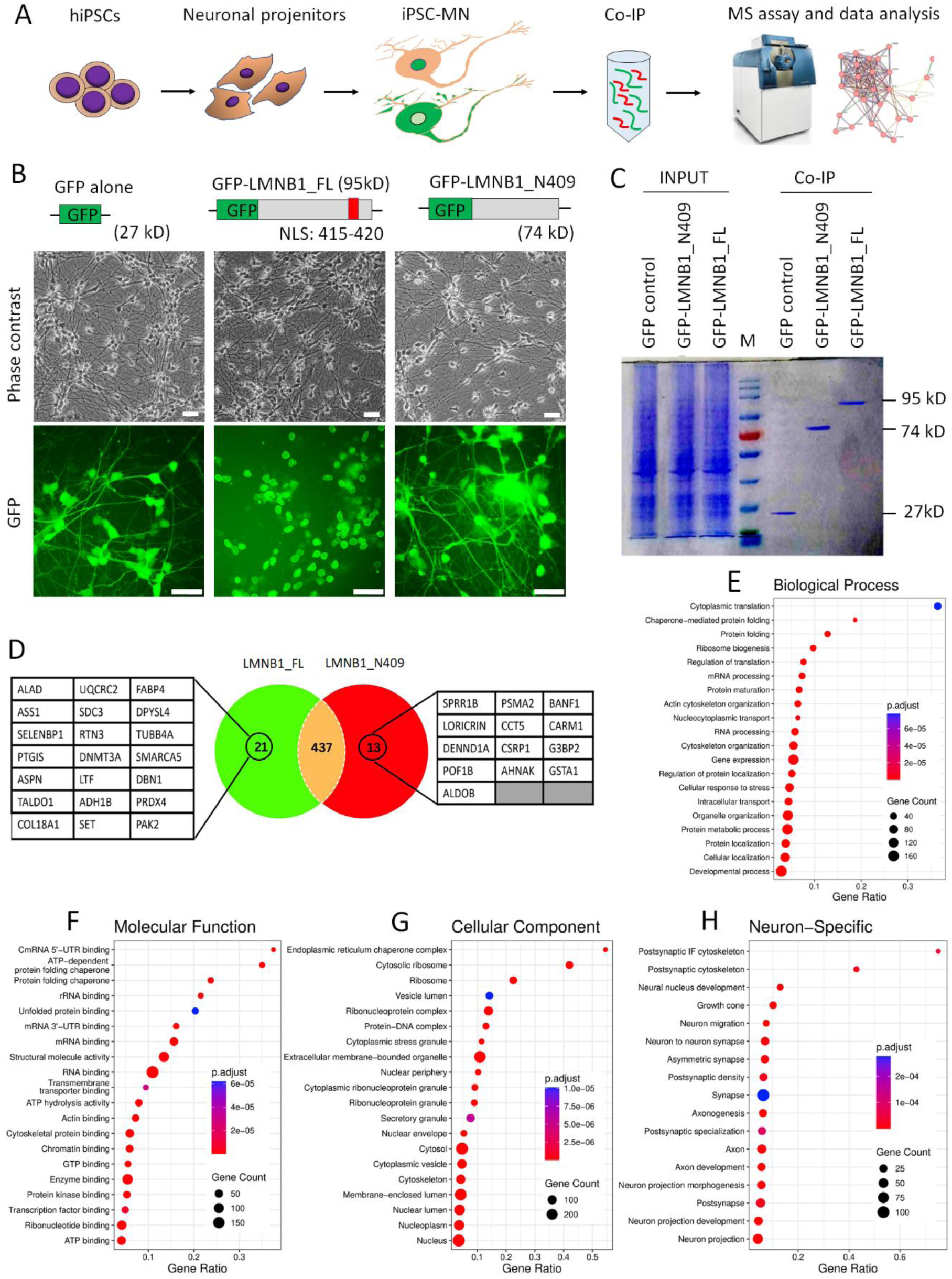
Identification of cytoplasmic Lamin B1-interacting proteins in hiPSC-derived motor neurons. (**A**) Schematic illustrating the workflow for identifying cytoplasmic Lamin B1-interacting proteins in human induced pluripotent stem cell-derived motor neurons (hiPSC-MNs) using co-immunoprecipitation followed by mass spectrometry (Co-IP/MS). (**B**) Representative fluorescence micrographs of hiPSC-MNs at 10 dpi, expressing GFP, full-length Lamin B1 with an N-terminal GFP tag (LMNB1_FL), or a truncated mutant lacking the NLS (LMNB1_N409). The truncated LMNB1 mislocalizes to the cytoplasm and neuronal processes. Scale bars: 50 µm. (**C**) Coomassie Brilliant Blue-stained SDS-PAGE gel of immunoprecipitated (IPed) samples from (B) using anti-GFP nanobody-conjugated with agarose beads. Input: 1% of total cell lysate; IP: 10% of IPed sample per lane. (**D**) Venn diagram showing the number of proteins bound to LMNB1_FL and/or LMNB1_N409. Unique interactors of LMNB1_FL and LMNB1_N409 are listed. (**E–H**) Gene Ontology (GO) analysis of the 471 total interactors identified from Co-IP MS, including 21 proteins exclusively associated with LMNB1_FL, 13 proteins exclusively associated with LMNB1_N409, and 437 shared interactors. (**E**) Biological Processes. (**F**) Molecular Functions. (**G**) Cellular Components. (**H**) Neuron-specific GO terms.

To enhance the efficiency of Co-IP, we ectopically expressed GFP-tagged nuclear Lamin B1 in healthy hiPSC-MNs. Comparing the impact of N-terminal and C-terminal tagged GFP on the subcellular distribution of nuclear Lamin B1, we observed that GFP fused at N-terminus exhibited normal nuclear distribution, while GFP fused at C-terminus dramatically changed the subcellular distribution of nuclear Lamin B1 (**Fig. S2A and B**). This alteration may be attributed to C-terminal tagged GFP disrupting the anchorage of nuclear Lamin B1 to the NE via farnesylation at its C-terminal cysteine residue ^2,6^. Consequently, we constructed lentiviral vectors expressing GFP alone, N-terminal GFP-tagged full length Lamin B1 (LMNB1_FL), and a truncated Lamin B1 (LMNB1_N409) lacking the NLS, which is expected to localize in the cytoplasm (**Fig. S2C**). Western blotting confirmed the expression of these proteins at the expected sizes (**Fig. S2D**). In addition to the NLS, protein Lamin B1 consists of a large N-terminal α-helical rod domain and a C-terminal Ig-like domain. Portions of these domains have been resolved with crystallography ^73,74^ (**Fig. S2E and F**). AlphaFold ^75,76^ predicted that the α-helical rod domain in LMNB1_N409 adopts a structure nearly identical to that of LMNB1_FL (**Fig. S2G**). This suggests that the truncated Lamin B1 can still form dimers and assemble into filaments.

Subsequently, we prepared hiPSC-MNs expressing GFP, GFP-LMNB1_FL, and LMNB1_N409 using our established protocols ^59,72,77^. As expected, the truncated Lamin B1 lacks the NLS, resulting in its mislocalization to the cytoplasm and neuronal processes (**Figs. 4B and S3A**). Whole cell extracts from these samples were used to identify cytosolic Lamin B1-associated proteins through Co-IP/MS. To enhance the efficiency and specificity of the Co-IP assay, we employed an anti-GFP Nanobody coupled with agarose beads to precipitate GFP and GFP tagged proteins. Coomassie Brilliant Blue stained SDS-PAGE verified robust expression of GFP and GFP tagged proteins in hiPSC-MNs, with each IPed sample showing a highly enriched single band at the expected size (**Fig. 4C**). This confirmed the efficiency and specificity of the Co-IP assay.

### Lamin B1-interacting proteins involved in numerous biological processes

Co-IP/MS data from hiPSC-MNs were processed based on two criteria. First, the abundance of GFP peptides in each group served as an internal loading control to normalize the abundance of identified proteins. Second, specific binding was defined by a threshold of at least 5-fold enrichment in GFP-LMNB1_FL or LMNB1_N409 compared to GFP controls. Consequently, we identified 471 proteins that bound to LMNB1_FL and/or LMNB1_N409, including 21 proteins that are exclusively bound to LMNB1-FL, 13 proteins that are exclusively bound to the truncated mutation LMNB1_N409, and 437 proteins that are bound to both (**Fig. 4D** and **Supplemental Table S4**). These targets include many proteins that have been reported to interact with human nuclear Lamin B1 (BioGRID:110187, [BioGRID | Database of Protein, Chemical, and Genetic Interactions](https://thebiogrid.org/)) ^78^, including nuclear lamina and nuclear envelope proteins (e.g., LMNA, LMNB2, TMEM201, and RPN1), nuclear and cytoplasmic structural proteins (e.g., ACTB, JUP, VIM, CKAP4, and MARCKS), chromatin and gene regulation proteins (e.g., CBX5, H2AC20, H2BC21, MACROH2A1, MACROH2A2, DDX39B, PARP1, and many heterogeneous nuclear ribonucleoproteins), cell cycle and signaling proteins (e.g., CTNNB1, CANX, and many heat shock proteins), and 14-3-3 proteins (e.g., YWHAB, YWHAE, and YWHAQ) ^79–83^ (**Supplemental Table S4**). The identification of these known Lamin B1-interacting proteins in iPSC-MNs further validates the efficacy and specificity of the Co-IP/MS analysis and supports the ability of truncated LMNB1_N409 to form filaments and interact with Lamin B1-associated proteins.

Gene Ontology (GO) analysis indicates that Lamin B1-interacting proteins in iPSC-MNs are highly enriched in biological processes related to cytoplasmic translation, gene expression, RNA processing, stability, and transport; protein folding, refolding, maturation and metabolism; cytoskeleton, organelle, and cellular component organization; and cellular responses to stress. (**Fig. 4E** and **Supplemental Table S5**). Their molecular functions primarily involve RNA, mRNA, GTP, and chromatin DNA binding; protein, unfolded protein, and cytoskeletal protein binding; structural molecule activity; ATP hydrolysis; signaling receptor activity; and G protein-coupled receptor activity; protein folding chaperone; and functions in structural constituent of synapse (**Fig. 4F**). Consistent with their functions, these proteins are highly enriched in cellular components such as membrane-bounded organelle, cytosolic ribosome, organelle lumen, ribonucleoprotein complex, vesicle, cytosol, and nucleoplasm (**Fig. 4G**). Remarkably, many GO terms are neuron-specific, such as postsynaptic cytoskeleton, neural nucleus development, growth cone, synapse, axon and axon development, neuron projection and neuron projection development (**Fig. 4H)**. Based on their functions and interactions, these Lamin B1-interacting proteins in human MNs mainly form six clusters, including peptide chain elongation, translation, and ribosome; cytoskeletal and focal adhesion; ATP synthesis and metabolism; mRNA processing; Chromatin assembly; and heat shock protein (HSP) family and protein folding (**Fig. S4**). These results support the notion that nuclear Lamin B1 plays critical roles in various biological processes, both general and neuron specific. Dysregulation of nuclear Lamin B1, either at expression level or subcellular distribution, may significantly impact cellular functions ^16,18,21^.

### Lamin B1-interacting proteins in hiPSC-MNs regulate NCT

Interestingly, many Lamin B1-interacting proteins identified in iPSC-MNs are critically involved in the regulation of nucleocytoplasmic transport (**Supplemental Table S6**). For example, KPNA6 (karyopherin subunit alpha 6, Importin-α family) and KPNB1 (Karyopherin Beta 1, Importin-β family) are essential for importing NLS-containing cargo into the nucleus ^84,85^. RAN (Ras-related Nuclear Protein), a small GTP binding protein, facilitates the translocation of RNA and proteins through the nuclear pore complex by maintaining the RAN-GTP/RAN-GDP gradient across the nuclear envelope ^70,85,86^. RANGAP1 (Ran GTPase Activating Protein 1) regulates the Ran GTPase cycle, which is crucial for both nuclear import and export ^85,87,88^. NPM1 (nucleophosmin 1), the most abundant nucleolar protein, acts as a chaperone for ribosomal proteins and core histones and shuttles between the nucleus and cytoplasm ^89^. Additionally, several heterogeneous nuclear ribonucleoproteins and RNA binding proteins (e.g., HNRNPA2B1, HNRNPC, HNRNPH1, HNRNPM, HNRNPU) are involved in RNA processing and regulate nuclear RNA transport ^90,91^. Numerous ribosomal proteins (e.g., RPL5, RPL11, RPL13, RPL15, RPL19, RPL23A, RPL27A, RPL35) are also participate in ribosome assembly and nuclear export ^92,93^. These results support our findings that dysregulation of nuclear Lamin B1 may directly or indirectly impair NCT activities, contributing to the nuclear accumulation of mRNAs and mislocalization of proteins (**Figs. 3**)^44^.

### Lamin B1-interacting proteins involved in neurodevelopment and neurological diseases

Additionally, we also identified more than 30 Lamin B1-interacting proteins that localize to neuron-specific structures, including synapses, axons, growth cones, neuron projections, and the postsynaptic density cytoskeleton. These proteins are involved in neurite outgrowth and axonal Transport (e.g., TUBB3, TUBB2A, TUBB2B, MAP1B, MAP2, MAP4, DPYSL2, GAP43, MARCKS, and ATP1A3), neurodevelopment and maturation (e.g., L1CAM, DPYSL2, DPYSL3, DPYSL5, GAP43, MAP1B, MAP2, MAP6, HNRNPU, and KHDRBS1), and synaptogenesis (e.g., SYN1, VAMP2, STXBP1, GDI1, GDI2, ELAVL2, ELAVL3, SFPQ, and RALY) (see detailed annotations in **Supplemental Table S7**). These results suggest that nuclear Lamin B1-interacting proteins are broadly involved in neurodevelopment, neuronal maturation, synaptogenesis, and overall neuronal functions. This conclusion is consistent with the critical roles of Lamin B1 in neurons as demonstrated in other model systems ^94–97^.

Remarkably, some of these Lamin B1-interacting proteins in iPSC-MNs are implicated in neurodegenerative diseases (e.g., FUS, HSPD1, HSPE1, SFPQ, and CTSD), neurodevelopmental disorders (e.g., TUBB3, STXBP1, MAP1B, MAP2, and KHDRBS1), and other neurological conditions (e.g., ATP1A3, L1CAM, and GDI1) (**Supplemental Table S7)**. For example, FUS (fused in sarcoma) protein is an RNA-binding protein whose mutations are linked to amyotrophic lateral sclerosis (ALS) and frontotemporal dementia (FTD) ^98–100^. SFPQ (splicing factor proline and glutamine rich) is an RNA-binding protein involved in pre-mRNA processing, and its dysfunction has been associated with several neurodegenerative diseases ^101,102^. CTSD (cathepsin D), a lysosomal protease, is implicated in Alzheimer’s disease and other neurodegenerative disorders when dysfunctional ^103^. STXBP1 (syntaxin binding protein 1), a core component of the presynaptic membrane-fusion machinery, is essential for neurotransmitter release. Mutations in this gene have been linked to neurodevelopmental disorders, including infantile epileptic encephalopathy and intellectual disability ^104,105^. KHDRBS1 (also known as Sam68; KH RNA binding domain containing, signal transduction associated 1) is a critical regulator of splicing for synaptic genes. Loss of Sam68 disrupts synaptic connectivity, impairs neuromuscular junction formation, and leads to the progressive degeneration of spinal motor neurons, resulting in motor coordination defects, ataxia, and abnormal social behaviors ^106–108^. Mutations in the ATP1A3 gene (ATPase Na+/K+ transporting subunit alpha 3) are associated with dystonia, epilepsy, and a range of other neurological disorders ^109,110^. The interaction of nuclear Lamin B1 with numerous proteins relevant to neurological diseases suggests that its dysregulation may represent a fundamental mechanism underlying various neuropathological conditions.

### Cytosolic mislocalized Lamin B1 sequesters factors critical for cellular processes and neuronal function

Statistical analysis of Co-IP/MS data from iPSC-MNs revealed that 41 proteins showed significantly altered binding to LMNB1_N409 compared to LMNB1_FL in iPSC-MNs. Among them, one protein (KPNA6) decreased binding, while 40 proteins displayed increased binding. Notably, the most highly enriched proteins associated with cytosolic Lamin B1 were 14-3-3 proteins, including YWHAB, YWHAG, and YWHAZ. (**Fig. 5A** and **Supplemental Table S8**). GO analysis demonstrated that cytosolic Lamin B1 broadly disrupts biological processes, including cellular metabolism, RNA processing and gene expression, protein translation, transport, and localization. It affects molecular functions such as RNA and protein binding, structural constituents of the ribosome, and cell adhesion. The dysregulated proteins are predominantly localized in the cytoplasm, including the extracellular region, cell junctions, cytosolic ribosomes, vesicles, and synapses (**Fig. 5B and Supplemental Table S8**). Network analysis further indicated that these proteins are mainly involved in protein metabolism, general metabolic pathways, the PI3K-Akt signaling pathway, and neuronal development (**Fig. 5C**).

**Figure 5.**
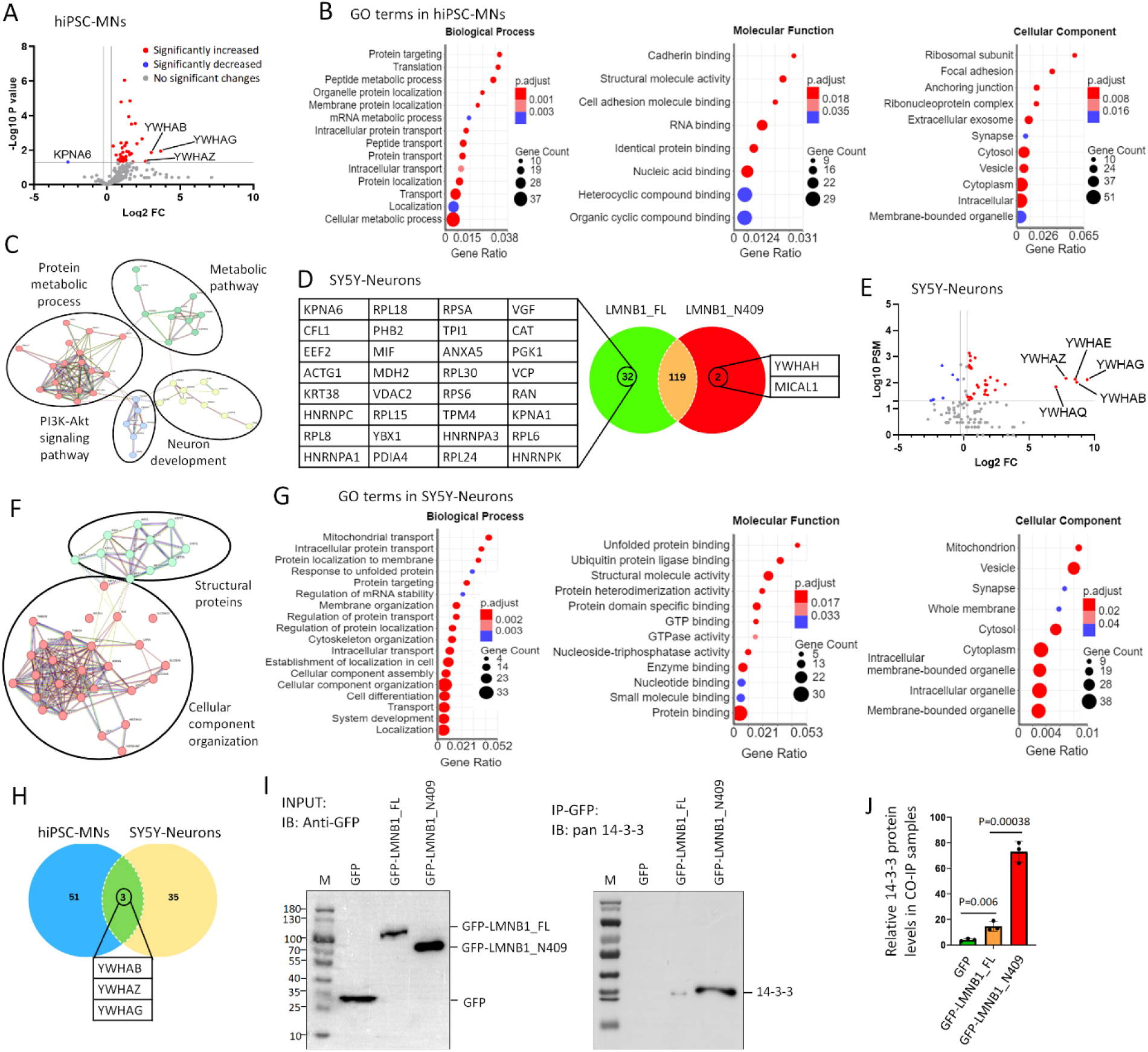
Cytoplasmic mislocalized Lamin B1 sequesters essential regulatory factors and strongly interacts with 14-3-3 proteins in human neurons. (**A**) Volcano plot illustrating differences in protein binding between LMNB1_FL and LMNB1_N409 in hiPSC-MNs. (**B**) Gene Ontology (GO) enrichment analysis of 54 proteins that exhibited significantly increased binding to LMNB1_N409 compared to LMNB1_FL, including 13 proteins uniquely associated with LMNB1_N409. Categories include Biological Process, Molecular Function, and Cellular Component. (**C**) Protein network analysis reveals that LMNB1_N409-enriched interactors are involved in protein metabolism, general metabolic pathways, PI3K-Akt signaling, and neuronal development. (**D**) Venn diagram showing the number of proteins bound to LMNB1_FL and/or LMNB1_N409 in SY5Y-derived neurons. Unique interactors of each form are listed. (**E**) Volcano plot of differential interactors between LMNB1_FL and LMNB1_N409 in SY5Y-neurons. (**F**) Network analysis shows that proteins with increased LMNB1_N409 binding in SY5Y-neurons cluster into two major functional categories: structural proteins and factors involved in cellular component organization. (**G**) GO analysis of 40 proteins with altered interaction with LMNB1_N409 in SY5Y-neurons, including 34 proteins with increased binding (highlighted in red in E) and 6 with decreased binding (blue in E). (**H**) Venn diagram summarizing proteins highly associated with LMNB1_N409 in both hiPSC-MNs and SY5Y-neurons. Three 14-3-3 isoforms are common interactors in both datasets. (**I**) Western blot validation of LMNB1 interactors. Input samples (2%) and Co-IP eluates from hiPSC-MNs expressing GFP, LMNB1_FL-GFP, or LMNB1_N409-GFP were probed using anti-GFP and pan–14-3-3 antibodies. M: protein marker. (**J**) Quantification of Co-IP 14-3-3 signals from (I). Data represent mean ± SEM from N = 3 independent experiments. Statistical significance was assessed using one-way ANOVA with Dunnett’s multiple comparisons test.

To determine whether cytosolic Lamin B1 similarly disrupts molecular interactions in non-MNs, we performed another Co-IP/MS analysis in neurons differentiated from human SH-SY5Y neuroblastoma cells (**Fig. S3B and C**). SY5Y cells can be induced to differentiate into neurons by treatment with retinoic acid (RA) ^44,111,112^. These differentiated neurons lack cholinergic markers, making them suitable for studying generic non-cholinergic neurons ^44^. After normalizing by GFP peptide levels, we identified 153 proteins that significantly interacted with LMNB1_FL and/or LMNB1_N409 in SY5Y-neurons. Among them, 32 proteins exclusively bound LMNB1_FL, 2 proteins exclusively bound LMNB1_N409 (YWHAH and MICAL1), and 119 proteins bound to both (**Fig. 5D**). Compared with LMNB1_FL, 40 proteins showed significantly altered binding to LMNB1_N409, including 34 with increased binding and 6 with decreased binding. (**Fig. 5E and Supplemental Table S8).** Compared to nuclear LMNB1_FL, the most significantly reduced binding proteins to cytosolic LMNB1_N409 are nuclear histones (e.g., HIST3H2A, HIST2H2BF, HIST1H4A) and nuclear Lamin A (LMNA), consistent with their subcellular distribution. Remarkably, 14-3-3 proteins (YWHAG, YWHAB, YWHAE, YWHAQ and YWHAZ) are the most highly enriched proteins associated with cytosolic Lamin B1 (**Fig. 5E).** Network analysis grouped these proteins into two major functional categories: cellular component organization and structural proteins (**Fig. 5F**). GO analysis showed that these proteins were significantly enriched in 76 biological processes and 28 molecular functions, many of which overlapped with findings in hiPSC-MNs, including protein transport and localization, mRNA processing, cytoskeletal organization, and synaptogenesis (**Fig. 5G and Supplemental Table S8**). Taking together, these results indicate that mislocalized nuclear Lamin B1 broadly interferes with essential biological processes and disrupts neuronal function in both induced human MNs and non-MNs.

### Cytosolic Lamin B1 strongly interacts with 14-3-3 proteins in induced human neurons

Interestingly, we found that 14-3-3 proteins are the most highly enriched group associated with cytosolic Lamin B1 in both Co-IP/MS datasets (**Fig. 5A and E**). The human 14-3-3 protein family comprises seven paralogs, namely β, γ, ε, η, σ, τ, and ζ (beta, gamma, epsilon, eta, sigma, tau and zeta) ^64,65,113^. Among these, six paralogs, including YWHAH (14-3-3 η), YWHAB (14-3-3β), YWHAG (14-3-3γ), YWHAZ (14-3-3ζ), YWHAE (14-3-3ε), and YWHAQ (14-3-3τ) were identified as the most abundant proteins associated with LMNB1_N409 in SY5Y-neurons (**Fig. 5D and E**). Notably, three of these paralogs (YWHAB, YWHAG and YWHAZ) were also identified in iPSC-MNs (**Figs. 5A**). These three paralogs were the only proteins found to exhibit significantly altered binding to LMNB1_N409 compared to LMNB1_FL in both neuron types (**Fig. 5H**).

To further validate the protein-protein interactions between cytosolic Lamin B1 and 14-3-3 proteins, we employed two complementary approaches. First, we performed Western blot analysis on Co-IP samples derived from iPSC-MNs expressing GFP, LMNB1_FL, or LMNB1_N409, as previously described. Using a pan-14-3-3 antibody, we observed a strong enrichment of 14-3-3 proteins in the LMNB1_N409 samples (**Fig. 5I and J**). In the second approach, we analyzed the colocalization of 14-3-3 proteins with cytosolic mislocalized Lamin B1 in reprogrammed DYT1 patient MNs. Consistent with our previous findings ^44^, MNs directly converted from DYT1 patient fibroblasts exhibited prominent cytoplasmic Lamin B1, whereas in healthy controls, Lamin B1 remained largely confined to the nucleus (**Fig. S5A**). Although the pan-14-3-3 antibody detects all paralogs expressed in both neurons and cocultured monolayer astrocytes, resulting in elevated background signal, it was clearly enriched and colocalized with mislocalized Lamin B1 in DYT1 neuronal processes (**Fig. S5B**). These results further support the colocalization and association between mislocalized nuclear Lamin B1 and 14-3-3 proteins in DYT1 MNs.

### The expressions of 14-3-3 paralogs vary across neurodevelopmental stages and are altered in DYT1 neurons

To determine whether 14-3-3 proteins are dysregulated in DYT1 neurons and to explore their potential roles in DYT1 pathogenesis, we first assessed the expression levels of their encoding genes using RT-PCR. Samples were collected from both iMNs and iGNs (glutamatergic neurons) derived from DYT1 iPSCs and isogenic controls at 8 and 16 days post-viral infection (dpi), a critical window for early neurodevelopment and neuronal maturation in cultured iPSC-derived neurons ^60–62,72,77^. The pan-neuronal markers *MAP2* exhibited robust expression and significant temporal upregulation (**Fig. 6B**), confirming the quality of the neuronal samples and the reliability of the RT-PCR assays. Consistent with previous reports showing that *YWHAS* (also known as *SFN* or Stratifin) is highly expressed in epithelial tissues and low in brain tissues ^114,115^, its transcript levels were barely detectable in induced neurons (**Fig. 6A**). Five paralogs (*YWHAG, YWHAE, YWHAH, YWHAQ*, *YWHAZ*) showed temporal upregulation in both GNs and MNs. In contrast, the *YWHAB* paralog exhibited temporal upregulation only in iGNs, but not in iMNs (**Fig. 6A and B**). These results suggest the differential expressions of 14-3-3 paralogs across developmental stages and neuronal subtypes. Notably, the paralogs *YWHAB, YWHAG,* and *YWHAE* were significantly downregulated in DYT1 MNs at 16 dpi compared to healthy controls (**Fig. 6B**), suggesting the dysregulation of these three paralogs during neuronal maturation in DYT1 MNs. Given that *YWHAB* and *YWHAG* are commonly identified as strongly associated with mislocalized Lamin B1 in both iPSC-MNs and SY5Y-neurons (**Fig. 5H**), we selected these two paralogs for further investigation to elucidate their roles in neurodevelopment and DYT1 pathogenesis.

**Figure 6.**
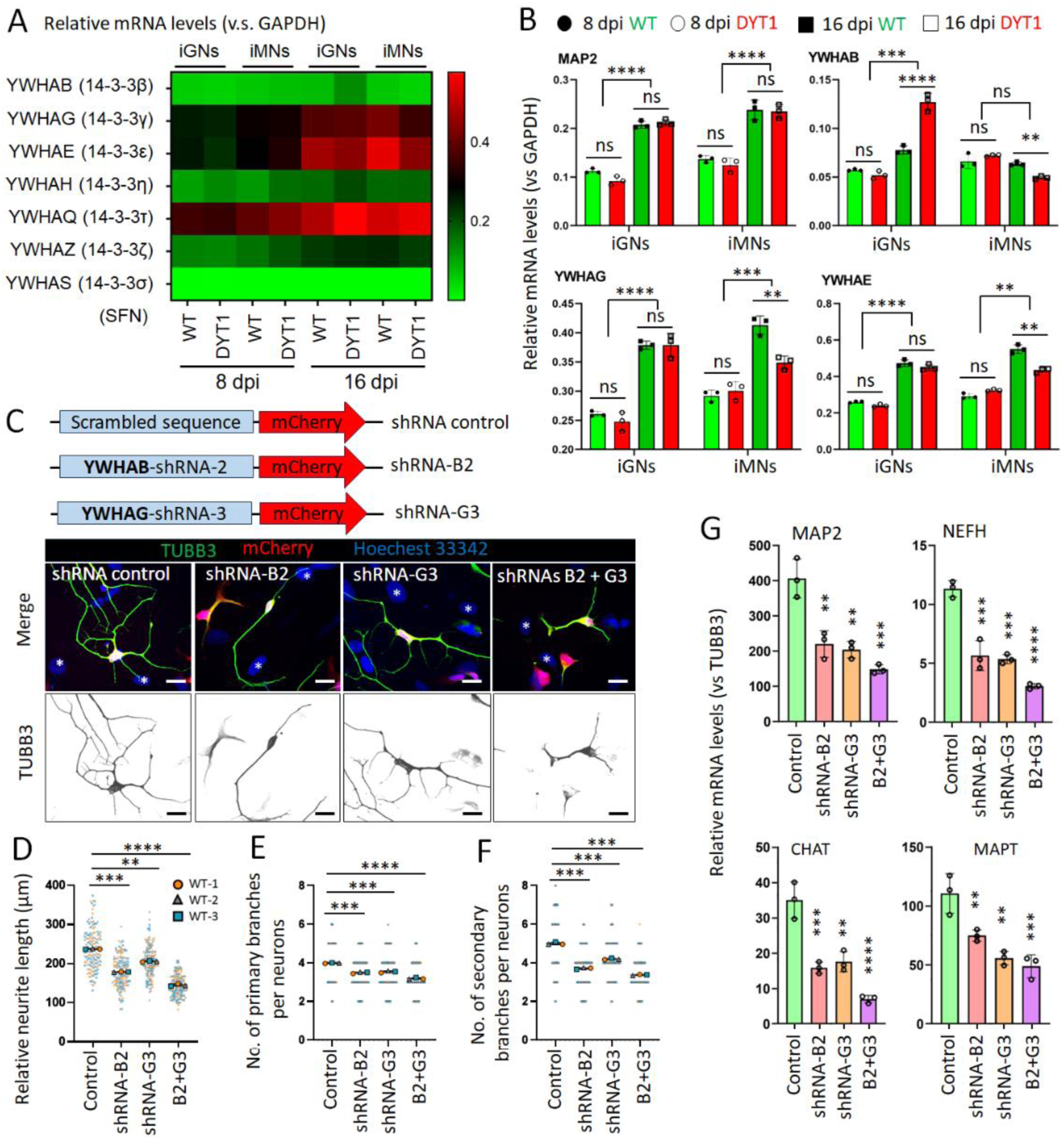
The expression of 14-3-3 proteins is altered in DYT1 neurons, and they are required for neurodevelopment and maturation. (**A**) A heatmap summarizes the gene expression of 14-3-3 paralogs in both WT (Control hiPSC-1) and DYT1 (DYT1 hiPSC-1) neurons at 8 and 16 dpi. Gene expression was analyzed by RT-PCR and normalized with GAPDH. iPSC-GNs, iPSC-derived glutamatergic neurons; iMNs, iPSC-derived motor neurons. SFN (Stratifin), the official name of the isoform YWHAS, the expression of which is extremely low in neurons. (**B**) Relative mRNA levels of indicated genes in both WT and DYT1 neurons (iGNs and iMNs) at 8 and 16 dpi. The plots show mean ± s.d. from three biological replicates (WT hiPSC-1, −2, and −3). ns, no significant difference; ** p<0.01; *** p<0.001; **** p<0.0001. One-way ANOVA with Dunnett’s multiple comparisons test. (**C**) Confocal micrographs of healthy iPSC-MNs at 10 dpi with transductions of lentiviruses expressing indicated shRNAs (mCherry +). Scale bars: 20 µm. (**D**) Relative neurite length of healthy iPSC-MNs at 10 dpi. N (objects) = 3. n (Neurons) = 50 for each object highlighted with distinct colors. Compared to control: ** p<0.01, *** p<0.001, **** p<0.0001. One-way ANOVA with Dunnett’s multiple comparison test. (**E**) Number of primary branches per neuron at 10 dpi. N (objects) = 3. n (Neurons) = 50 for each object highlighted with distinct colors. Compared to control: *** p<0.001, **** p<0.0001. One-way ANOVA with Dunnett’s multiple comparison test. (**F**) Number of secondary branches per neuron at 10 dpi. N (objects) = 3. n (Neurons) = 50 for each object highlighted with distinct colors. Compared to control: *** p<0.001. One-way ANOVA with Dunnett’s multiple comparison test. (**G**) Relative mRNA levels of indicated genes in healthy iPSC-MNs at 10 dpi. The plots show mean ± s.d. from three biological replicates (WT hiPSC-1, −2, and −3). Compared to control: ** p<0.01; *** p<0.001; **** p<0.0001. One-way ANOVA with Dunnett’s multiple comparison test.

### Downregulation of 14-3-3 proteins impairs neurodevelopment and maturation

To manipulate the gene expression of the *YWHAB* and *YWHAG*, we constructed lentiviral vectors for either overexpression (with GFP-tagged) or downregulation (with mCherry reporter). Each target gene was knocked down using three different shRNAs, each directed to a distinct site within the coding region (**Fig. S6**). Due to the lack of isotype-specific antibodies for detecting endogenous protein levels, we performed co-transfection of GFP-tagged 14-3-3 constructs along with their corresponding shRNAs to validate overexpression and knockdown efficiency based on GFP fluorescent signal intensity (**Fig. S6A-D**) and ectopically expressed protein levels (**Fig. S6E and F**). Through this screening, *YWHAB*-shRNA-2 (B2) and *YWHAG*-shRNA-3 (G3) were identified as the most effective constructs for downregulating *YWHAB* and *YWHAG*, respectively. We prepared lentiviruses expressing these shRNAs to transduce healthy iPSC-MNs. A vector containing a scrambled sequence served as the control (**Fig. 6C**). Strikingly, MNs transduced with these shRNAs exhibited significantly shorter neurite lengths and fewer primary and secondary branches (**Fig. 6C-F**), suggesting that 14-3-3β and 14-3-3γ are important for neurite outgrowth at early neurodevelopment. Consistently, RT-PCR analysis indicated that the expression levels of genes involved in neurodevelopment and maturation significantly decreased in iPSC-MNs with the downregulation of 14-3-3β and/or 14-3-3γ compared to controls, including MAP2, NEFH (Neurofilament Heavy Chain), CHAT (Choline Acetyltransferase) and MAPT (Microtubule-Associated Protein Tau). (**Fig. 6G**). Notably, their effects appeared additive or synergistic, as MNs co-transduced with both shRNAs showed more pronounced phenotypes than those transduced with either shRNA alone (**Fig. 6C-G**).

### Overexpression of 14-3-3 proteins rescues DYT1 neurite outgrowth and maturation

Since 14-3-3 proteins are among the most abundant in the brain and are involved in numerous biological processes essential for neuronal function ^64,65^, we propose that the impaired neurodevelopment of DYT1 neurons may result from functional disruption or reduced availability of 14-3-3 proteins due to their sequestration by cytoplasmic Lamin B1. Therefore, overexpression of 14-3-3 proteins may help rescue the cellular deficits observed in DYT1 neurons. We generated lentiviral vectors expressing GFP-tagged YWHAB or YWHAG, and the vector expressing GFP alone works as a control. The density of seeded DYT1 iPSC-MNs and the transduction efficiency are similar across conditions (**Fig. S7A**). Transduced neurons have been examined from 7 to 21 dpi. Compared to GFP controls or YWHAB and YWHAG transduction negative (GFP-) neurons, DYT1 iPSC-MNs with the overexpression of YWHAB or YWHAG (GFP+) showed significantly longer neurite length and more primary and secondary branch numbers at both 7 dpi (**Fig. S7B-E**) and 14 dpi (**Fig. 7A-D**). Consistently, the expression levels of genes involved in neurodevelopment and maturation significantly increased in DYT1 iPSC-MNs with the upregulation of YWHAB or YWHAG compared to GFP controls (**Fig. S7F**). These results suggest that the elevation of 14-3-3β or 14-3-3γ protein levels can rescue the impaired neurite outgrowth and maturation in DYT1 neurons at early developmental stages.

**Figure 7.**
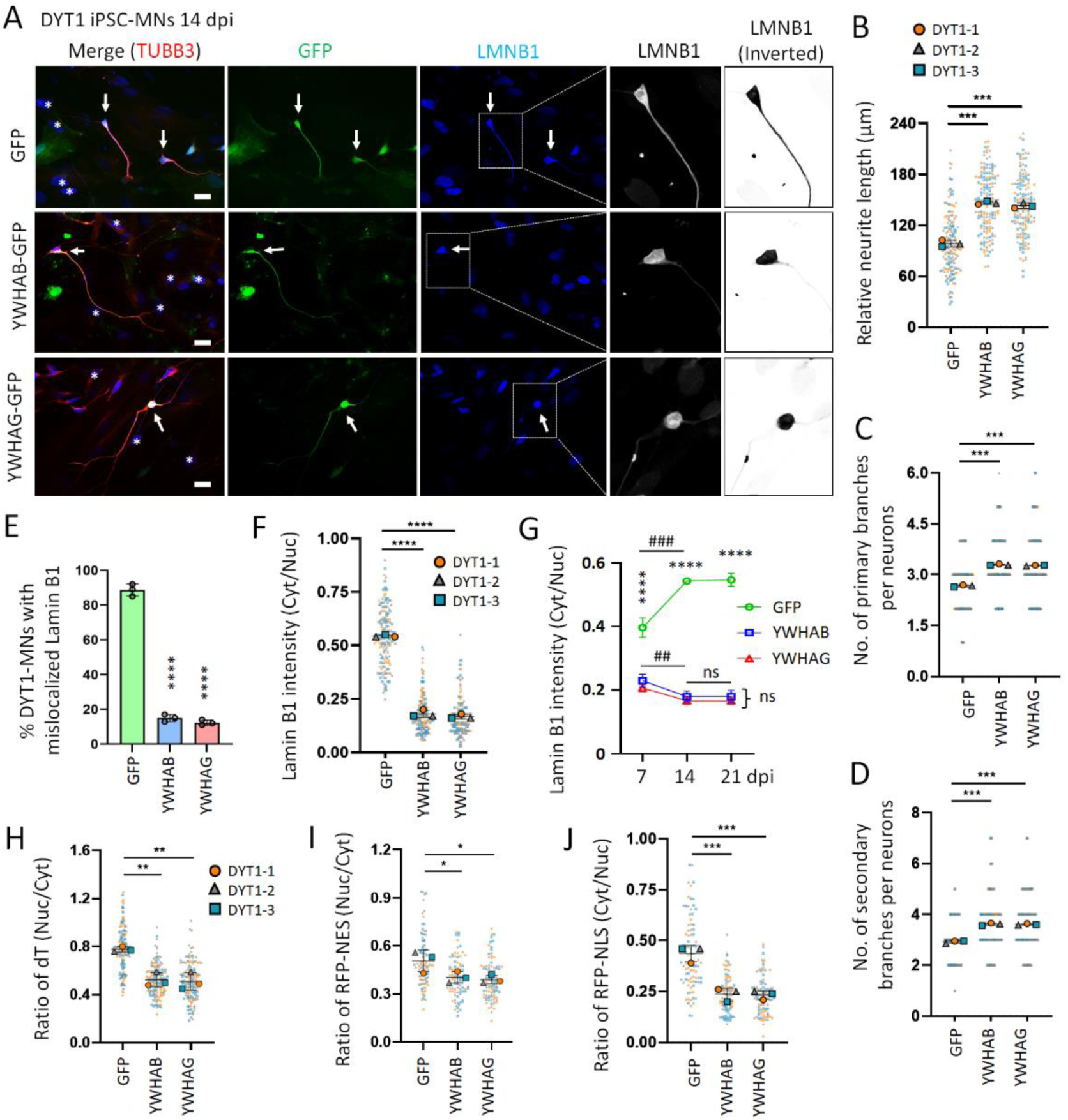
Overexpression of 14-3-3 proteins in DYT1 neurons mitigates Lamin B1 mislocalization and rescues neuronal deficits. (**A**) Fluorescence micrographs of DYT1 iPSC-MNs (DYT1 hiPSC-1) at 14 dpi with overexpression of GFP-tagged 14-3-3 proteins. LMNB1 signals in the highlighted neurons are also shown at higher magnification in grayscale. Arrows indicate transduction positive (GFP +) neurons, and asterisks mark cocultured astrocytes. Scale bars: 20 µm. (**B**) Relative neurite length of DYT1 iPSC-MNs at 14 dpi. N (objects) = 3 (DYT1 hiPSC-1, −2, AND −3). n (Neurons) = 50 for each object highlighted with distinct colors. Compared to GFP control: *** p<0.001. One-way ANOVA with Dunnett’s multiple comparison test. (**C**) Quantification of primary branches per neuron. N (objects) = 3. n (Neurons) = 50 for each object highlighted with distinct colors. Compared to control: *** p<0.001. One-way ANOVA with Dunnett’s multiple comparison test. (**D**) Quantification of secondary branches per neuron. N (objects) = 3. n (Neurons) = 50 for each object highlighted with distinct colors. Compared to control: *** p<0.001. One-way ANOVA with Dunnett’s multiple comparison test. (**E**) Quantification of the percentage of DYT1 MNs exhibiting mislocalized Lamin B1 at 14 dpi, corresponding to panel A. Each group includes 150 neurons from three biological replicates. (**F**) Quantification of the cytoplasmic-to-nuclear Lamin B1 ratio in DYT1 iPSC-MNs at 14 dpi, corresponding to images shown in panel A. N (objects) = 3. n (Neurons) = 50 for each object highlighted with distinct colors. Compared to control: **** p<0.0001. One-way ANOVA with Dunnett’s multiple comparison test. (**G**) Dynamic changes in the cytoplasmic-to-nuclear Lamin B1 ratio of DYT1 iPSC-MNs from 7 to 21 dpi. N (objects) = 3. The value of each condition at each time point calculated from 50 neurons. Compared to GFP control: *** p<0.001; **** p<0.0001. One-way ANOVA with Dunnett’s multiple comparison test. Compared to 7 dpi within each group: ns, no significant difference; ## p<0.01, ### p<0.001. One-way ANOVA with Dunnett’s multiple comparison test. (**H**) Quantification of fluorescence in situ hybridization (FISH) assay in DYT1 iPSC-MNs at 14 dpi, shown as the ratio of nuclear-to-cytoplasmic oligo-dT signal intensity. N (objects) = 3. n (Neurons) = 50 for each object highlighted with distinct colors. Compared to GFP control: ** p<0.01. One-way ANOVA with Dunnett’s multiple comparison test. (**I**) Quantification of protein export assay using RFP-NES reporter in DYT1 iPSC-MNs at 14 dpi, shown as the ratio of nuclear to cytoplasmic RFP signal intensity. N (objects) = 3. n (Neurons) = 50 for each object highlighted with distinct colors. Compared to GFP control: * p<0.05. One-way ANOVA with Dunnett’s multiple comparison test. (**J**) Quantification of protein import assay using RFP-NLS reporter in DYT1 iPSC-MNs at 14 dpi, shown as the ratio of cytoplasmic to nuclear RFP signal intensity. N (objects) = 3. n (Neurons) = 50 for each object highlighted with distinct colors. Compared to GFP control: *** P<0.001. One-way ANOVA with Dunnett’s multiple comparison test.

### Overexpression of 14-3-3 proteins mitigates Lamin B1 mislocalization and rescues impaired NCT in DYT1 neurons

To investigate whether upregulation of 14-3-3 proteins affects Lamin B1 subcellular distribution, we co-stained TUBB3 and nuclear Lamin B1 in DYT1 iPSC-MNs transduced with lentiviruses expressing GFP-tagged YWHAB or YWHAG (**Fig. 7A**). Consistent with previous findings (**Fig. S5**) ^44^, in the GFP control, the majority (∼85%) of DYT1 neurons exhibited marked mislocalization of Lamin B1 to the cytoplasm and neuronal processes. In sharp contrast, this frequency dropped to less than 20% in neurons overexpressing YWHAB or YWHAG at 14 dpi (**Fig. 7E**). Correspondingly, the average of cytoplasmic-to-nuclear ratio of Lamin B1 signal intensity significantly decreased from above 0.5 in GFP controls to approximately 0.2 in 14-3-3-overexpressing neurons at 14 dpi (**Fig. 7F**). In DYT1 iPSC-MNs, cytoplasmic mislocalization of Lamin B1 became evident at 7 dpi and progressively increased, with the cytoplasmic-to-nuclear ratio peaking at 21 dpi, a time point when iPSC-MNs reach maturation ^44,62^ (**Fig. 7G**). Strikingly, DYT1 MNs overexpressing 14-3-3 proteins showed markedly reduced cytoplasmic Lamin B1 signal as early as 7 dpi and maintained lower cytoplasmic-to-nuclear ratios through at least 21 dpi, with no significant difference between paralogs 14-3-3β and 14-3-3γ (**Fig. 7G**). These results indicate that elevated levels of 14-3-3 proteins mitigate Lamin B1 mislocalization in DYT1 neurons, and that the paralogs 14-3-3β and 14-3-3γ are similarly effective.

Given that Lamin B1 mislocalization could be a major contributor to cellular deficits in DYT1 neurons, including impaired nuclear transport ^44,62^, we asked whether mitigating Lamin B1 mislocalization through overexpression of 14-3-3 proteins could also rescue impaired NCT. To test this, we first examined nuclear mRNA export by a FISH assay with oligo-dT probes, as previously described (**Fig. 3**) ^68,70^. DYT1 MNs overexpressing 14-3-3 proteins showed a significantly reduced nuclear-to-cytoplasmic dT signal ratio compared to GFP controls (**Fig. 7H**), indicating that upregulation of 14-3-3 proteins improved the efficiency of nuclear mRNA export. Next, we assessed protein nuclear transport in DYT1 MNs by co-transducing lentiviruses expressing GFP-tagged 14-3-3 proteins together with RFP-NES or RFP-NLS reporters. Only GFP and RFP double-positive neurons were analyzed. The nuclear-to-cytoplasmic ratio of RFP-NES was used to assess nuclear export, while the cytoplasmic-to-nuclear ratio of RFP-NLS was used to assess nuclear import, as described previously^62,68,70^. In DYT1 MNs overexpressing 14-3-3 proteins, both ratios were significantly reduced compared to GFP controls, indicating improved efficiencies of both protein nuclear export and import (**Fig. 7I and J**). Notably, the reduction in the RFP-NES ratio was moderate (**Fig. 7I**), whereas the decrease in the RFP-NLS ratio was more pronounced (**Fig. 7J**), suggesting that 14-3-3 proteins may enhance protein nuclear import more effectively than export.

## DISCUSSION

In Eukaryotic cells, nuclear morphology and nuclear transport play critical roles in the maintenance of cellular homeostasis, particularly important for neuronal functions due to their specialized cellular morphology and structure ^84,85,116^ (**Fig. S8A**). The deformed neuronal nucleus in the DYT1 mouse model was first observed two decades ago ^53^, and subsequent studies have reported similar abnormalities in different model systems carrying Torsin mutations ^43,44,47,49^. However, the precise mechanisms by which the ΔE mutation leads to nuclear deformation and DYT1 neuronal deficits have remained elusive. In our study, we have demonstrated that dysregulation of nuclear Lamin B1, in terms of both expression level and subcellular localization, is a distinct cellular deficit in DYT1 cells, including DYT1 patient fibroblasts and reprogrammed patient-derived neurons. Additionally, we provide evidence that upregulation of nuclear Lamin B1 results in a thickened nuclear lamina and causes the deformation of the nucleus and the impaired NCT (**Fig. S8B**). Importantly, our proteomic studies using induced human neurons have revealed that mislocalized nuclear Lamin B1 extensively interferes with critical factors and signaling pathways, leading to widespread cellular dysfunction in affected neurons. Remarkably, 14-3-3 proteins are the strongest interactions with cytoplasmic Lamin B1 (**Fig. S8B**). Downregulation of 14-3-3 proteins significantly disrupt neurodevelopment and maturation in induced human neurons, while upregulation of 14-3-3 proteins can rescue DYT1 neuronal deficits, at least in part, by mitigating the Lamin B1 mislocalization (**Fig. S8C**). These findings shed light on the molecular mechanisms underlying the deformed nucleus and cellular dysfunction in DYT1 cells. Moreover, our results contribute to a deeper understanding of the pathophysiology of DYT1 dystonia and provide new evidence regarding the nuclear Lamin B1 dysregulation in human diseases.

In mammals, most differentiated somatic cells contain both A-type and B-type Lamin proteins. Intriguingly, we observed that only nuclear Lamin B1 was dysregulated in DYT1 cells, while Lamin A/C remained unaffected. This finding suggests the existence of distinct spatial organization and/or functional regulation between Lamin B1 and Lamin A/C. Consistently, recent studies have demonstrated differential localization and molecular architecture of Lamin B1 and Lamin A/C within the nuclear lamina ^1,117^. Lamin B1 forms an outer concentric ring of the nuclear lamina and is located closer to the NE compared with Lamin A/C ^117^. Compared with wild-type Torsin, the mutated ΔE protein exhibits a distinct subcellular distribution pattern, with high enrichment around the NE ^35,118^. Thus, the more peripheral localization of Lamin B1 may make it more directly associated with the NE and more susceptible to alterations caused by NE-associated proteins, such as the mutated Torsin ΔE. Accumulating evidence indicates that mutations in the *LMNB1* gene or fluctuations in its expression can lead to abnormal nuclear morphology, chromatin organization and gene expression formation ^6,119^. These abnormalities have been associated with disrupted embryonic development and organogenesis of the brain ^120,121^, as well as the onset of various diseases, including cancers and neurological disorders ^17,19,122^. The dysregulation of nuclear Lamin B1 in DYT1 cells could be associated with changes in gene expression, potentially related to the reorganization of chromatin structure induced by nucleus deformation ^123,124^. Further investigation is required to fully understand the detailed mechanisms underlying the specific dysregulation of nuclear Lamin B1 in DYT1 cells.

In addtion to the expression levels, the subcellular localization of Lamin B1 is also critical for maintaining cell functions. Our research has demonstrated that nuclear Lamin B1 mislocalization is a unique cellular deficit in DYT1 neurons. Lamin B1 progressively accumulates in the cytoplasm and neuronal processes during the differentiation process, peaking at the maturation stage ^44^. In this study, we further demonstrated that the mislocalized nuclear Lamin B1 extensively disrupts critical factors and signaling pathways in induced human neurons. The disrupted biological processes include cellular metabolism, cytoskeleton organization, synaptogenesis, protein and RNA transport and localization, resulting in widespread cellular dysfunction. More interestingly, using iPSC-MNs, we identified that cytoplasmic Lamin B1 disrupts multiple factors critical for nuclear transport, neurite outgrowth, neurodevelopment, and neuronal maturation. These findings support our hypothesis that Lamin B1 dysregulation is a key pathogenic factor in DYT1 dystonia. Specifically, nuclear deformation and the sequestration of essential molecular components by mislocalized Lamin B1 appear to be major mechanisms contributing to neuronal deficits in DYT1 (**Fig. S8B**).

One of the most exciting findings in this study is that 14-3-3 proteins are the most prominently affected targets disrupted by mislocalized Lamin B1. These proteins are highliy expressed in the brain and play imporatnt roles in CNS development, particularly in neurodevelopmental processes ^64,65^. Dysregulaton of 14-3-3 proteins has been implicated in various neurological disorders, including Parkinson’s disease, Alzheimer’s disease, and several neurodevelomental disorders ^65,114,125,126^. Consistent with this, our study demonstated that downregulation of 14-3-3 proteins significantly impaired neurite outgrowth and neuronal maturation in iPSC-derived MNs. Intrestingly, we also observed decreased expression of some 14-3-3 paralogous (β, γ and ε) in DYT1 neurons. We propose that sequesteration of 14-3-3 proteins by cytoplasmic Lamin B1 in DYT1 neurons may substantially interfere with their physiological functions, ultimately leading to neuronal dysfunction (**Fig. S8B**). This may represent a key molecular mechanism by which mislocalized Lamin B1 contributes to the pathogenesis of DYT1 dystonia.

Excitingly, we also demonstarted that overexpression of 14-3-3 can largely rescue DYT1-associated neuronal deficits, including disrupted neurite outgrowth and impaired nuclear transport, most likely by mitigating Lamin B1 mislocalization. As molecular chaperones in either homodimer, heterodimer or monomer ^64,127,128^, 14-3-3 proteins may alleviate these deficits through multiple mechanisms. First, elevated levels of 14-3-3 proteins may sequester cytoplasmic Lamin B1 via liquid-liquid phase separation (LLPS) ^129^, thereby preventing it from interfering with other essential neuronal factors (**Fig. S8C**). Second, since 14-3-3 proteins contain NLS, they may facilitate proper trafficking and localization of their binding partners ^130,131^. This includes promoting the nuclear import of mislocalized Lamin B1 in DYT1 neurons, thereby reducing cytoplasmic Lamin B1 levels and maintaining a lower cytoplasmic-to-nuclear ratio, as observed in DYT1 iPSC-MNs with overexpression of 14-3-3. Third, binding of 14-3-3 proteins can influence the activity and stability of their targets ^132,133^, potentially promoting Lamin B1 degradation through the ubiquitin-proteasome system. Additionally, 14-3-3 proteins may regulate transcription of genes involved in neurodevelopment and maturation^134^, thereby further contributing to the rescue of DYT1 neuronal deficits. Further investigation into the molecular mechanisms by which 14-3-3 proteins exert these effects will advance our understanding of DYT1 dystonia and aid in the development of therapeutic strategies.

We acknowledge that several aspects of our investigation and analysis have limitations. First, we currently have limited insight into the molecular mechanisms underlying the increased interaction between cytoplasmic Lamin B1 and 14-3-3 proteins. One possibility is that the altered subcellular distribution of Lamin B1 enhances its interactions with specific 14-3-3 paralogs that predominantly localize in the cytoplasm. Given that 14-3-3 proteins are known to bind their targets in a phosphorylation-dependent manner ^128,135^, another potential mechanism involves post-translational modifications of Lamin B1, such as phosphorylation, that may influence its binding affinity. This represents an important area for future investigation. In this study, patient-derived MNs were used as a primary model system, based on growing evidence that dysfunction of spinal neural circuits, particularly spinal cord MNs, contributes significantly to the pathophysiology of DYT1 dystonia ^44,51,136^. However, dystonia is increasingly recognized as a network disorder involving multiple brain regions, including the basal ganglia, cerebellum, and sensorimotor cortex ^137–146^. Whether the molecular changes observed in MNs can be recapitulated in other neuronal subtypes remains to be determined. Given that the dysregulated factors (e.g., LMNB1, 14-3-3 proteins) and pathways (e.g., nuclear transport, synaptogenesis, neurodevelopment) identified in DYT1 MNs are broadly conserved across cell types, we propose that similar mechanisms may be shared by other neurons affected in DYT1 dystonia, potentially contributing to dysfunction of the neuronal network. Expanding the research in these directions will provide a more comprehensive understanding of the molecular mechanisms underlying DYT1 dystonia and may inform the development of potential therapeutic strategies for this debilitating neurological disorder.

## ACKNOWLEDGMENTS

We thank members of the Ding laboratory (Dr. Masuma Akter, Mr. Casey A. Coutee, and Mr. Md. Kobirul Islem) for their technical assistance and Dr. William T. Dauer (University of Texas Southwestern) for discussions. We also acknowledge the support of the Shared Instrumentation Facility at the Louisiana State University for the electron microscopy (EM) analysis; the Mass Spectrometry Core Facility and Redox Molecular Signaling Core of Centers of Biomedical Research Excellence at LSU Health Sciences Center Shreveport for the LC-MS/MS analysis; and LSU Health Shreveport Research Core Facilities for confocal imaging and RT-PCR assay. This work was supported by National Institutes of Health (NIH) (NS133252 to B.D.).

## AUTHOR CONTRIBUTIONS

B.D. conceptualized and designed experiments. B.D., Y.X., and J.S. performed EM study. Y.D., B.D., M.S., and X.S. performed proteomic analyses. Y.D., B.D., and M.A.H. performed cell culture, imaging and quantification. H.C. performed AlphaFold predication. B.D., Y.D. and M.S. analyzed and interpreted data. B.D., Y.D., and M.S. prepared figures. B.D. wrote the original manuscript. B.D., Y.D., C.L.Z. and Y.L. edited the manuscript. All authors have read and provided inputs to the manuscript.

## DECLARATION OF INTERESTS

The authors declare no competing interests.

## METHODS

### Cell lines and culture condition

The use of human cell lines, including human induced pluripotent stem cells (hiPSCs), in this study was approved by the Institutional Review Board (IRB) of Louisiana State University Health Sciences Center at Shreveport (LSUHSC-S, IRB ID: STUDY00002945). All cell lines used in this study were listed in **Supplemental Table S1**. HEK 293T cells (CRL-11268) and SH-SY5Y (CRL-11266) cells were purchased from ATCC. DYT1 fibroblast cell lines and sex- and age-matched healthy controls were obtained from the *Coriell Institute for Medical Research* or *NINDS Human Cell and Data Repository* (NHCDR). Human induced pluripotent stem cell (hiPSC) lines have been generated in our previous studies ^44,60,61^. All hiPSCs were maintained in complete mTeSR1 medium (STEMCELL Technologies) on Matrigel (Corning) coated dishes at 37 °C and 5% CO_2_ with saturating humanity, and the medium was daily replaced.

The recipes of cell culture medium were:

1. HEK medium: DMEM (Gibco) supplemented with 10% fetal bovine serum (FBS, Corning, NY, USA) and 1% penicillin/streptomycin (ThermoFisher).
2. Fibroblast medium: DMEM supplemented with 15% FBS and 1% penicillin/streptomycin.
3. hiPSC medium: complete mTeSR1 medium and 1% penicillin/streptomycin.
4. Neuronal induction medium: DMEM: F12: neurobasal (2:2:1), 1% N2 (Invitrogen), 1% B27 (Invitrogen), 1% penicillin/streptomycin, and supplemented with 10 mM Forskolin (FSK, Sigma-Aldrich), 1 mM dorsomorphin (DM, Millipore, MA, United States) and 10 ng/mL basic fibroblast growth factor (bFGF, PeproTech).
5. Neuronal maturation medium: DMEM: F12: neurobasal (2:2:1), 1% N2 (Invitrogen), 1% B27 (Invitrogen), 1% penicillin/streptomycin, and supplemented with 5 mM FSK and 10 ng/mL each of BDNF, GDNF, and NT3 (PeproTech).
6. KOSR medium: DMEM/F12 medium (Gibco) with 20% KnockOut Serum Replacement (KOSR, ThermoFisher), 1% GlutaMax (Gibco), 1% non-essential amino acid (NEAA, (Gibco), 50 μM β-mercaptoethanol (β-ME, Gibco), 1% penicillin/streptomycin and 10 ng/mL bFGF.
7. Neurosphere medium (NSP medium): DMEM/F12 medium containing 1% N2, 1% GlutaMax, 1% NEAA, 50 µM β-ME, 1% penicillin/streptomycin, 8 μg/mL Heparin, 20 ng/mL bFGF and 20 ng/ml epidermal growth factor (EGF, PeproTech).
8. Neural progenitor cell (NPC) medium: DMEM/F12 and neurobasal medium (1:1) containing 0.5% N2, 1% B27, 1% GlutaMax, 1% NEAA, 50 µM β-ME, 1% penicillin/streptomycin, 10 ng/mL EGF and 10 ng/mL bFGF.
9. SH-SY5Y medium: DMEM supplemented with 15% FBS and 1% penicillin/streptomycin.
10. SH-SY5Y differentiation medium: DMEM supplemented with 3% FBS, 10 µM all-trans-Retinoic acid (Sigma) and 1% penicillin/streptomycin.

### Plasmid construction and lentivirus production

A third-generation lentiviral vector (*pCSC-SP-PW-IRES-GFP*) was used to express reprogramming factors NEUROG2-IRES-GFP-T2A-SOX11, NEUROG2-IRES-SOX11, ISL1-T2A-LHX3, and NGN2-IRES-ISL1-T2A-LHX3 as previously reports ^44,62,72,147^. The same lentiviral vector was used to express GFP-tagged LMNB1 and its truncated mutation (LMNB1_N409), GFP-tagged YWHAB and YWHAG. PCR amplified GFP sequence with incorporated BamHI (5’) and NheI (3’) sites, and amplified cDNA sequences of full-length LMNB1, LMNB1_N409, YWHAB and YWHAG with NheI (5’) and XhoI (3’) restriction sites. Inserts of GFP and gene of interest were ligated into the lentiviral vector between the BamHI and XhoI restriction sites. Another third-generation lentiviral vector (*LV-CAG-mCherry-miRE-Luc*) was used to express shRNAs targeting LMNB1, YWHAB and YWHAG as described in previously reports ^44,62,148^. Briefly, shRNA oligos targeting the gene coding regions were synthesized, PCR amplified with flanking miR-30 sequences, and ligated into the above lentiviral vector at the *XhoI* and *EcoRI* restriction sites. The targeting sequences (gRNAs) were listed in **Supplemental Table S2.** The dual reporter 2Gi2R was generously provided by Dr. Fred H. Gage ^69^. The transport reporters of RFP-NES and RFP-NLS were constructed as previously reported ^62^.

All plasmids were verified by restriction enzyme digestions and DNA sequencing. Each of these vectors was co-transfected with packaging plasmids (Addgene #12251, #12253, and #12259) into HEK293T cells for virus production ^149,150^. Replication-incompetent lentiviruses were produced and viral supernatants were collected at 48 hrs and 72 hrs post-transfection as previously described ^58,151^. The viral supernatants were filtered through 0.45 μm syringe filters and stored at 4 °C prior to cell transduction.

### Direct conversion of adult fibroblast cells into MNs

Directly converted MNs were prepared via lentiviral delivery of 4 factors (*NEUROG2*, *Sox11*, *ISL1* and *LHX3*) as previously described ^44,58,147^. In brief, fibroblast cells were plated at a density of 1×10^4^ cells/cm^2^ onto Matrigel-coated dishes. Cells were transduced the next day with lentiviral supernatants supplemented with 6 mg/mL polybrene. Fibroblast medium was refreshed after overnight incubation. One day later, the culture was replaced with neuronal induction medium and half-changed every other day until 14 days post-viral infection (dpi). A replating procedure was used to purify induced neurons ^44,58^. The purified cells were cultured in neuronal maturation medium onto Matrigel-coated coverslips with or without the presentence of astrocytes depending on desired experiments. The medium was half changed twice a week until analysis.

### Generation of hiPSC-derived MNs

MNs were prepared from hiPSCs as previously described ^57,59,72^. Briefly, hiPSCs were cultured in mTeSR1 medium with 10 µM all-trans-retinoic acid (RA, Sigma) and 0.5 mM VPA in Matrigel-coated 6-well plates for 7 days. Cells were then digested with Versene and gently pipetted into small clumps supplemented with 10 μM Y-27632 (STEMCELL Technologies). Cell clumps were cultured in KOSR medium for 4 days, followed by cultured in NSP medium for another week. The neurospheres were then dissociated into single cells with Accutase (Innovative Cell Technologies) and maintained in neural progenitor cell medium. For MN differentiation ^60,72,77^, neural progenitor cells were plated into Matrigel-coated plates at a density of 3×10^4^ cells/cm^2^ and transduced with a lentiviruse expressing *NEUROG2-IRES-ISL1-T2A-LHX3*. Culture medium was replaced the next day with neuronal maturation medium. Neurons were dissociated with Accutase on day 5 and replated onto Matrigel-coated coverslips with or without the presentence of astrocytes depending on desired experiments. The medium was half changed twice a week until analysis.

### Generation of SH-SY5Y-derived neurons

The SH-SY5Y neuroblastoma cells were cultured on gelatin-coated plates and maintained in SH-SY5Y medium. To induce neuronal differentiation, the media was switched to SH-SY5Y differentiation medium. The medium was half changed every other day. SY5Y cells gradually became neuron-like morphology within one week. To ectopically overexpress genes in SY5Y-derived neurons, transduced lentiviruses expressing genes of interest when cells reached about 50% confluence. After an overnight exposure to lentivirus, the media was replaced the next day with SH-SY5Y differentiation medium, and half changed twice a week until analysis.

### Immunocytochemistry (ICC)

Cultured cells at the indicated time points were fixed with 4% paraformaldehyde (PFA) in PBS for 15 min at room temperature (RT) and then permeabilized and blocked for 1 hr in blocking buffer (PBS containing 0.2% Triton X-100 and 3% BSA). They were subsequently incubated overnight with primary antibodies in blocking buffer at 4 °C, followed by washing and incubation with corresponding fluorophore-conjugated secondary antibodies. Cell nuclei were counterstained with Hoechst 33342 (HST, ThermoFisher). Primary antibodies used in this study were listed in **Supplemental Table S3**.

### Fluorescent in situ hybridization (FISH)

FISH was performed as described previously ^62,68,70^. Briefly, cultured cells were washed once with PBS, and fixed with 4% PFA for 30 min at RT. Cells were permeabilized with 0.2% TritonX-100 for 10 min, followed by two washes with PBS. Samples were equilibrated with hybridization buffer composed of 2 X SSC, 10% dextran sulfate, 10 mM ribonucleoside-vanadyl complex (RVC; New England Biolabs) and 20% formamide. A mixture of digoxigenin (DIG*)-*labeled oligo-dT or dA probes (0.2 ng/µL) and yeast tRNA (0.2 µg/µL) were heated to 95 °C for 5 min and immediately chilled on ice. Probes were then combined with equal volumes of 2 X hybridization buffer and incubated with samples overnight at 37 °C. Anti-DIG antibody (Sigma, 11333089001, 1:100) and corresponding fluorophore-conjugated secondary antibodies (ThermoFisher) were used to detect oligo-dT signals. Anti-MAP2 (Abcam, ab5392, 1:10000) antibody and HST were used to determine neuronal soma and nuclei, respectively. All reagents and solutions were prepared in nuclease-free water.

### Protein nuclear transport assay

Protein nuclear transport was analyzed using a dual reporter (2Gi2R) or individual reporters (RFP-NES, RFP-NLS) as previously described ^62,68,70^. Briefly, cultured cells were transduced with a lentivirus expressing reporters. For fibroblast cells, the reporter lentivirus was transduced when cell growth reached about 50% confluency. The signal density and distribution of GFP and RFP were analyzed at 7 dpi. For iPSC-derived neurons, reporter lentiviruses were co-transduced with lentiviruses expressing MN reprogramming factors. The signal densities and distributions of GFP and/or RFP were analyzed at the indicated time points.

### Imaging and quantification analysis

Images were obtained with Nikon A1R, Leica TCS SP5, Zeiss LSM700 (63×/1.4 NA) confocal microscopes or an inverted fluorescence microscope (Olympus, CKX53). For nuclear morphology assay, we zoomed in on individual nucleus by using the ImageJ software and defined the abnormal nucleus based on two criteria: 1) at least one obvious sharp angle or a deep invagination, and 2) at least half area of the nucleus losing the smooth outline. Signals of nuclei (HST) or nuclear Lamins were used to distinguish the nucleus and cytoplasm. For fluorescence intensity quantification, as large an area as possible was measured within the nucleus or cytoplasm as previously described ^68,70^. An unbiased approach for data collection was employed. The person who analyzed the images was completely blinded to the sample information. The “mean” values were used to quantify an average signal intensity.

### Transmission electron microscopy (TEM) and immunogold labeling

Cells on Matrigel-coated glass coverslips were fixed overnight at 4 °C with a solution composed of 4% paraformaldehyde, 1% glutaraldehyde, 1.5 µM MgCl_2_ and 5 µM CaCl_2_ in 0.1 M cacodylate buffer (pH7.4). Samples were then washed twice with cacodylate buffer, treated for 1.5 hr at RT with 1% osmium tetraoxide (OsO_4_, EMS) and 0.8% K3Fe(CN6) in cacodylate buffer, and prestained with 2% uranyl acetate in water for 2 hr in the dark at RT. The subsequent steps of dehydration, infiltration, and embedding were the same as previously described ^44,152,153^. The resin blocks were polymerized overnight at 60 °C and sectioned with a diamond knife (Diatome) on a Leica Ultracut UCT 6 ultramicrotome (Leica Microsystems). Sections were post stained with 2% uranyl acetate in water and lead citrate. Images were acquired on a JEOL JEM 1200EX transmission electron microscope equipped with a tungsten source at 120kV using a Morada SIS camera (Olympus). The electron-dense areas immediately beneath the INM were measured with ImageJ (NIH) and the ratio of electron-dense area to the nuclear perimeter was used to evaluate the thickness of nuclear lamina.

For immunogold labeling, a different fixative solution (composed of 4% paraformaldehyde and 0.1% glutaraldehyde) was used to preserve antigens. The sectioned grids were incubated with block solution (5% BSA in PBS buffer) to minimize non-specific labeling and transferred to drops of diluted primary antibody in 2% BSA in 0.05% PBST buffer. The primary antibody is Rabbit anti-LMNB1 (Proteintech, 12987-1-AP, 1:100) and the secondary antibody was Goat anti-Rabbit IgG conjugated with 15 nm gold particles (Cytodiagnostics, AC-15-01-05, 1:100). Samples were treated without primary antibody served as negative controls. EM micrographs were taken at both low and high magnifications with the focus on NE structure. To examine the subcellular distribution of nuclear Lamin B1, gold particles were counted and grouped into three regions, nucleoplasm, nuclear envelope, and cytoplasm.

### Co-immunoprecipitation coupled with mass spectrometry (Co-IP/MS) assay

Cultured cells were harvested and lysed using a lysis buffer containing 50 mM Tris-HCl (pH 7.4), 150 mM NaCl, 0.5 mM EDTANa2, 2% Triton-X 100, 1mM PMSF, and DNase I and RNase A. The lysate was centrifuged (15000 rpm, 30 min, 4 °C) to remove any cellular debris. The supernatant was transferred to new tubes and used for Co-IP. Anti-GFP Nanobody coupled with agarose beads (Proteintech, gta) was used to precipitate GFP and GFP tagged proteins as previously described ^154–156^. Briefly, the beads were equilibrated using wash buffer 1 (50 mM Tris-HCl (pH 7.4), 150 mM NaCl, 0.5 mM EDTANa2, and 2% Triton-X 100) and blocked with 1% BSA in wash buffer 1 for 30 min at 4 °C. And then the beads were resuspended in the lysate and incubated on a rotator for at least 2 h at 4 °C. After spinning down, the beads were washed with wash buffer 1 and then with wash buffer 2 (50 mM Tris-HCl (pH 7.4), 500 mM NaCl, 0.5 mM EDTANa2, and 2% Triton-X 100) followed by resuspension in elution buffer (50 mM Tris-HCl (pH 8.5), 1% glycerol, 1% SDS) and boiling for 10 min.

For LC-MS/MS analysis of these eluted proteins, 10 µg of protein from each sample was loaded into 12% SDS-PAGE, and then stained with Imperial Protein Stain (ThermoFisher). The Gel lanes of each sample were cut into 2mm X 2mm cubes, and then performed trypsin digestion followed by disulfide bond reduction with dithiothreitol and cysteine alkylation with iodoacetamide, according to the manufacturer’s protocol of the Thermo Scientific™ Gel Tryptic Digestion kit. The digested peptides were vacuum-dried and resolved in the LC/MS grade water with 0.1% (v/v) formic acid, followed by untargeted discovery proteomics analysis. This LC-MS/MS analysis was carried out using an Ultimate 3000 RSLCnano system connected to an Orbitrap Exploris 480 mass spectrometer. The digested peptides (5.0 µL) were loaded onto a trap column (PepMap C18, 2 cm × 100 μm, 100 Å) at a flow rate of 20 μL/min using 0.1% formic acid, and separated on an analytical column (EasySpray 50 cm × 75 μm, C18 1.9 μm, 100 Å) with a flow rate of 300 nL/min with a linear gradient of 5 to 45% solvent B (100% ACN, 0.1% formic acid) over a 120 min gradient. Both precursor and fragment ions were acquired in the Orbitrap mass analyzer. Precursor ions were acquired in m/z range of 375-1500 with a resolution of 120,000 (at m/z 200). Precursor fragmentation was carried out using the higher-energy collisional dissociation method using normalized collision energy (NCE) of 32. The fragment ions were acquired at a resolution of 150,000 (at m/z 200). The scans were arranged in top-speed method with 3 sec cycle time between MS and MS/MS. Ion transfer capillary voltage were maintained at 2.1 kV.

The raw Mass Spec data were analyzed using Proteome Discover (version 2.5, Thermo Fisher Scientific) software package with SequestHT using species-specific fasta database and the Percolator peptide validator. Cysteine alkylation was set as a fixed modification, and deamidation of asparagine residues was selected as variable modifications. The abundance of GFP peptides in each group (GFP, GFP-LMNB1_FL, and LMNB1_N409) served as an internal loading control to normalize the abundance of identified proteins. Specific binding was defined by a threshold of at least 5-fold enrichment in GFP-LMNB1_FL or LMNB1_N409 compared to GFP controls. For iPSC-MNs, proteins showing significant changes in binding to the truncated mutant were selected based on a ≥2-fold change compared to the full-length control and a p-value < 0.05 across three replicates. For SH-SY5Y neurons, protein abundance changes were evaluated based on fold-change alone, as only one replicate was used for verification purposes. To acquire protein-protein interaction networks and their functional clusters, network analysis was performed using an online database (https://string-db.org) and Gene Ontology (GO) analysis was conducted using PANTHER (protein annotation through evolutionary relationship) classification system (http://www.pantherdb.org/) ^157^.

### Western blotting analysis

Cells were lysed in lysis buffer composed of 50 mM Tris-HCl buffer (pH 8.0), 150 mM NaCl, 1% NP40, 1% Triton X-100, 0.1% SDS, 0.5% sodium deoxycholate, and protease inhibitor cocktail (Roche). Equal amounts of cell lysates (20 µg per lane) were used for SDS-PAGE and western blot analysis as previously described ^44,156,158^. Primary antibodies and dilutions were listed in **Supplemental Table S3.** HRP-conjugated secondary antibodies and Clarity Western ECL substrate (Bio-Rad) were used to visualize the protein bands with a BioRad ChemiDoc.

### Quantitative real-time PCR analysis

As described previously ^62,159^, total RNA was extracted from cultured neurons using TRIzol (Life Technologies). cDNA synthesis reactions were performed using 0.5 µg of total RNA from each sample with the SuperScriptIII First-Strand kit (Life Technologies) and random hexamer primers. Real-time PCR was performed in triplicate using primers, SYBR GreenER SuperMix (Invitrogen), and the BIO-RAD CFX-96 Fast Real-Time PCR system. Target mRNA levels were normalized to the reference gene *GAPDH* or *TUBB3* by 2^-ΔΔCt^ method. Sequences of RT-PCR primers were listed in **Supplemental Table S2.**

### Protein structure prediction using AlphaFold

Protein structures of full-length Lamin B1 and its truncated mutant were predicted using AlphaFold3 via the Colab Multimer notebook with default settings [75,76]. Briefly, the amino acid sequences of human full-length Lamin B1 and its truncated form were obtained from UniProt and submitted to the AlphaFold3 server for structure prediction. The predicted models were analyzed using PyMOL, and those with the highest confidence scores were selected for further analysis. For protein superposition, structural models of full-length LMNB1 (residues 1– 586) and its truncated form (residues 1–409) were generated using AlphaFold3. Root-mean-square deviation (RMSD) values for Coil1 and Coil2 domains were calculated in PyMOL after structural alignment of the two models.

### Statistical analysis

Statistical analysis was done in GraphPad Prism. The D’Agostino & Pearson omnibus normality test was conducted first to determine if the data are normally distributed. If the data passed the normality test, one-way or two-way ANOVA was used to determine significance. If the data did not pass the normality test, the Kruskal-Wallis test was used to determine significance. Results are expressed as mean ± SD of at least three biological replicates or three independent experiments, and *P* < 0.05 is considered significant.

## SUPPLEMENTARY INFORMATION

### Supplementary Figures and Figure Legends

**Figure S1.**
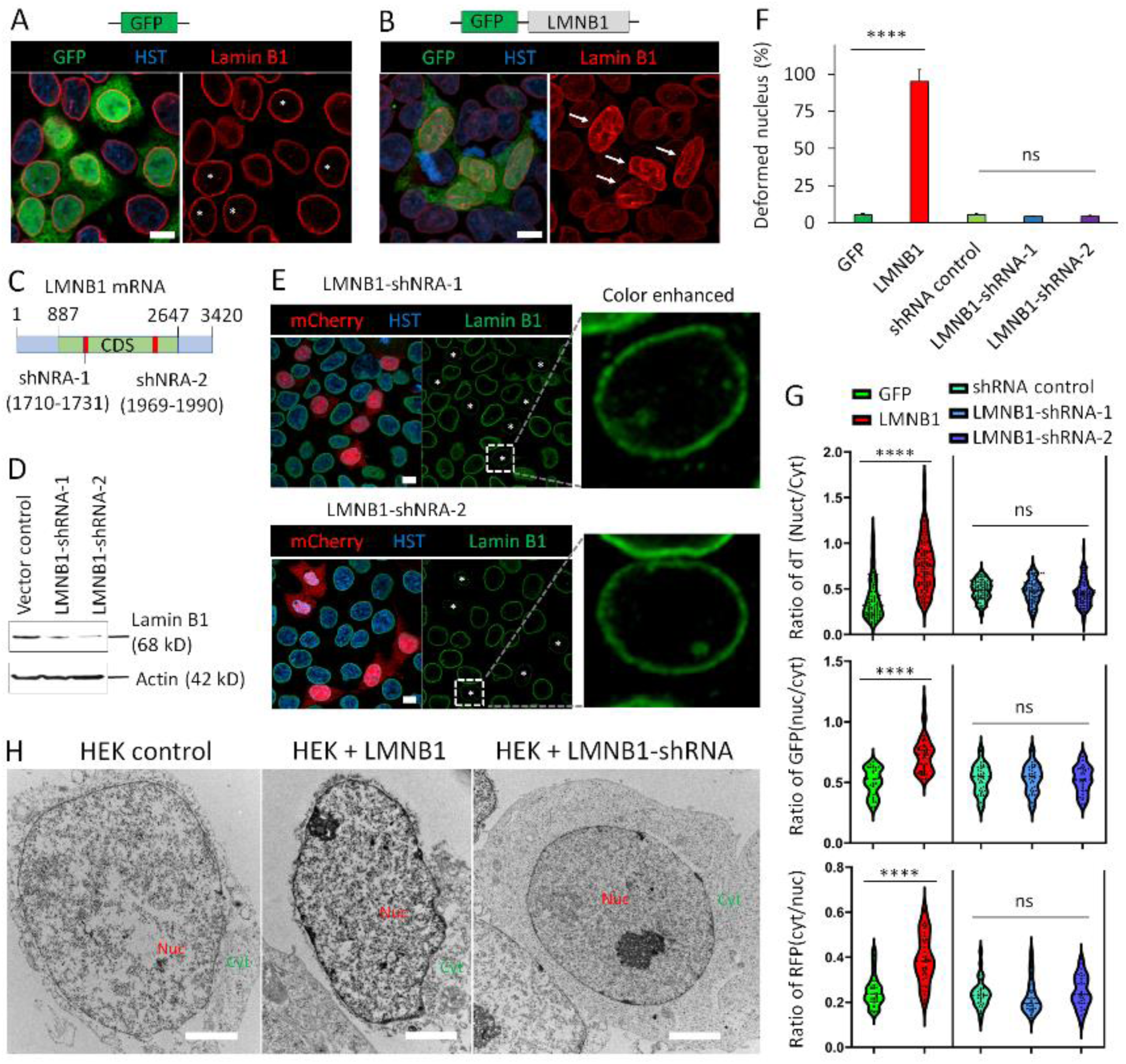
Upregulation of nuclear Lamin B1 significantly disrupts nuclear envelope morphology and nucleocytoplasmic transport, whereas downregulation does not. (**A**) Representative confocal micrograph of human embryonic kidney (HEK) cells ectopically express GFP. Asterisks indicate typical nuclear morphology of GFP-positive cells. Scale bar: 10 µm. (**B**) Representative confocal micrograph of HEK cells ectopically express nuclear Lamin B1 and the GFP reporter. Arrows indicate deformed nucleus with overexpression of Lamin B1. Scale bar: 10 µm. (**C**) A schematic shows two shRNAs targeting different sites within the coding sequence (CDS) of the LMNB1 transcript. (**D**) Western blot analysis shows that both LMNB1-targeting shRNAs efficiently down-regulate nuclear Lamin B1 protein levels. Beta-actin serves as a loading control. (**E**) Representative confocal images of HEK cells expressing LMNB1-shRNAs. The transfected cells express mCherry and are marked with asterisks in Lamin B1 staining. Cells with white squares are also shown at higher magnification with enhanced contrast to highlight nuclear envelope morphology. Scale bars: 10 µm. (**F**) Quantification of HEK cells with nuclear deformation under indicated conditions. n (cells) > 100 from three triplicates. ns, no significant difference; **** p<0.0001. Student’s t-test. (**G**) Quantification of nucleocytoplasmic transport under indicated conditions. FISH assay, n (cells) > 100 from triplicates. Reporter assay, n (cells) > 50 from triplicates. ns, no significant difference; **** p<0.0001. Student’s t-test and one-way ANOVA. (**H**) Representative transmission electron microscopy (TEM) images of HEK nuclei under indicated conditions. Nuc, nucleus; Cyt, cytoplasm. Scale bars: 2 µm.

**Figure S2.**
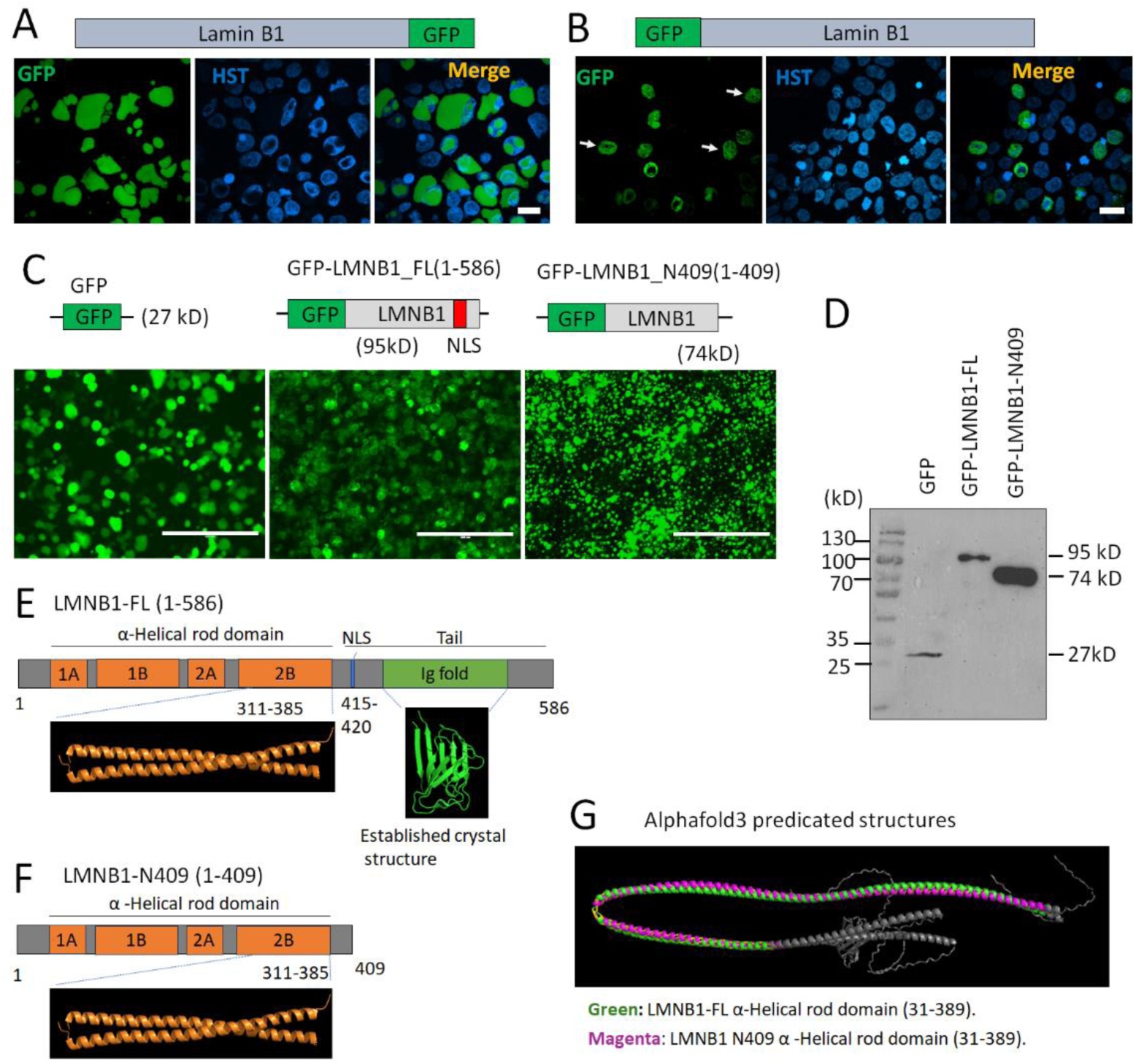
Overexpression of GFP-tagged full-length nuclear and truncated Lamin B1. (**A**) Micrograph of HEK cells expressing full-length Lamin B1 with a C-terminal GFP tag. Scale bar: 20 µm. (**B**) Micrograph of HEK cells expressing full-length Lamin B1 with an N-terminal GFP tag. Arrows indicate overexpressed LMNB1-GFP. Scale bar: 20 µm. (**C**) Micrographs of HEK cells expressing GFP alone, N-terminally tagged full-length Lamin B1 (LMNB1_FL), or a truncated Lamin B1 lacking the nuclear localization signal (LMNB1_N409). Scale bars: 200 µm. (**D**) Western blot showing expression of GFP, GFP tagged LMNB1_FL and LMNB1_N409, each at their expected molecular weight. (**E**) Schematic of full-length LMNB1 protein domains, including regions with known crystal structures: the α-helical rod domain 2B and the Ig-fold domain in the C-terminal tail. (**F**) Schematic of the LMNB1 truncation mutant (residues 1–409), highlighting the retained α-helical rod domain 2B and the absence of the C-terminal tail. (**G**) AlphaFold3-predicted structural comparison of full-length LMNB1 and its truncation mutant.

**Figure S3.**
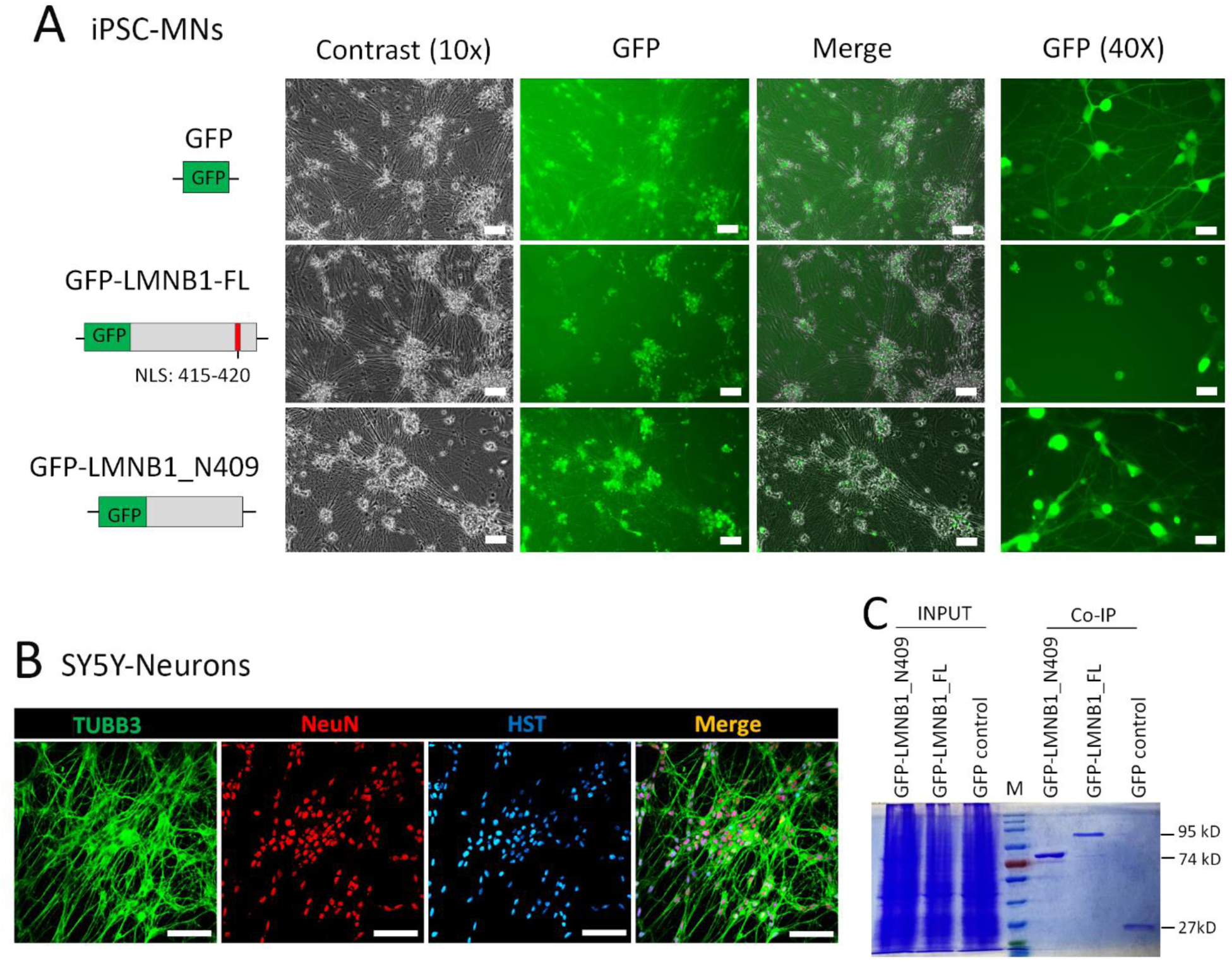
Overexpression of GFP-tagged Lamin B1 constructs in human induced neurons and immunoprecipitation sample preparation for proteomic analysis. (**A**) Micrographs of hiPSC-derived motor neurons at 10 days post-infection (dpi) with lentiviruses expressing GFP, GFP-tagged LMNB1_FL, or LMNB1_N409. The truncation mutant lacks the NLS and mislocalizes to the cytoplasm and neuronal processes. Scale bars: 100 µm (10×), 20 µm (40×). (**B**) Micrographs of neurons differentiated from the SH-SY5Y neuroblastoma cell line. Scale bar: 100 µm. (**C**) Coomassie Brilliant Blue-stained SDS-PAGE gel of immunoprecipitated (IP) samples prepared from SH-SY5Y-derived neurons using anti-GFP nanobody-conjugated magnetic agarose beads. Each lane was loaded with either 1% of total cell lysate (INPUT) or 10% of the IP sample.

**Figure S4.**
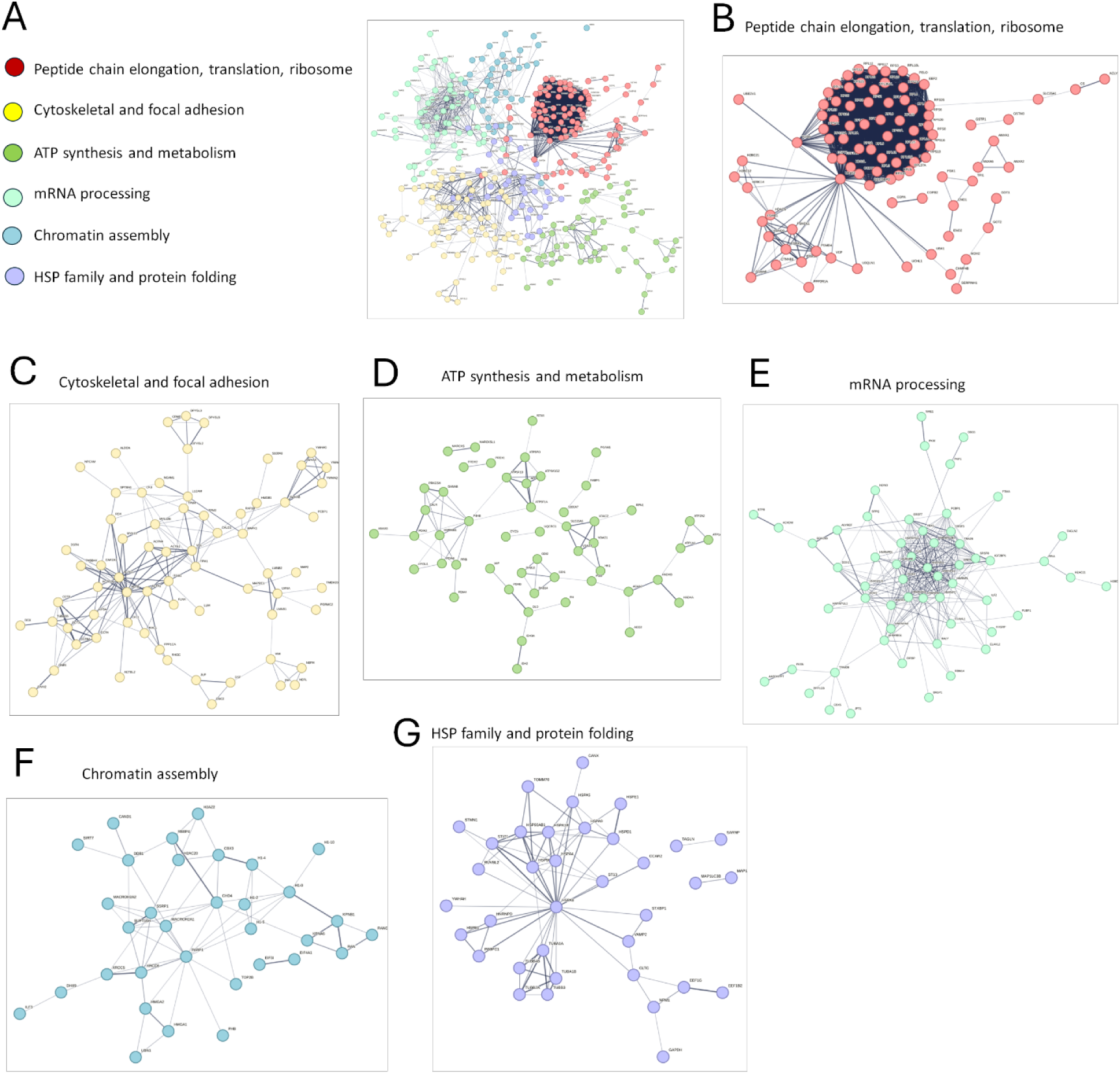
Lamin B1-interacting proteins in hiPSC-MNs are highly enriched in six clusters. (**A**) Network analysis reveals that Lamin B1-interacting proteins form six distinct clusters based on functional enrichment. (B–G) Detailed views of individual clusters: (**B**) Peptide chain elongation, translation, and ribosome-related proteins. (**C**) Cytoskeletal organization and focal adhesion components. (**D**) ATP synthesis and metabolic processes. (**E**) mRNA processing factors. (**F**) Chromatin assembly-related proteins. (**G**) Heat shock proteins (HSPs) and protein folding machinery.

**Figure S5.**
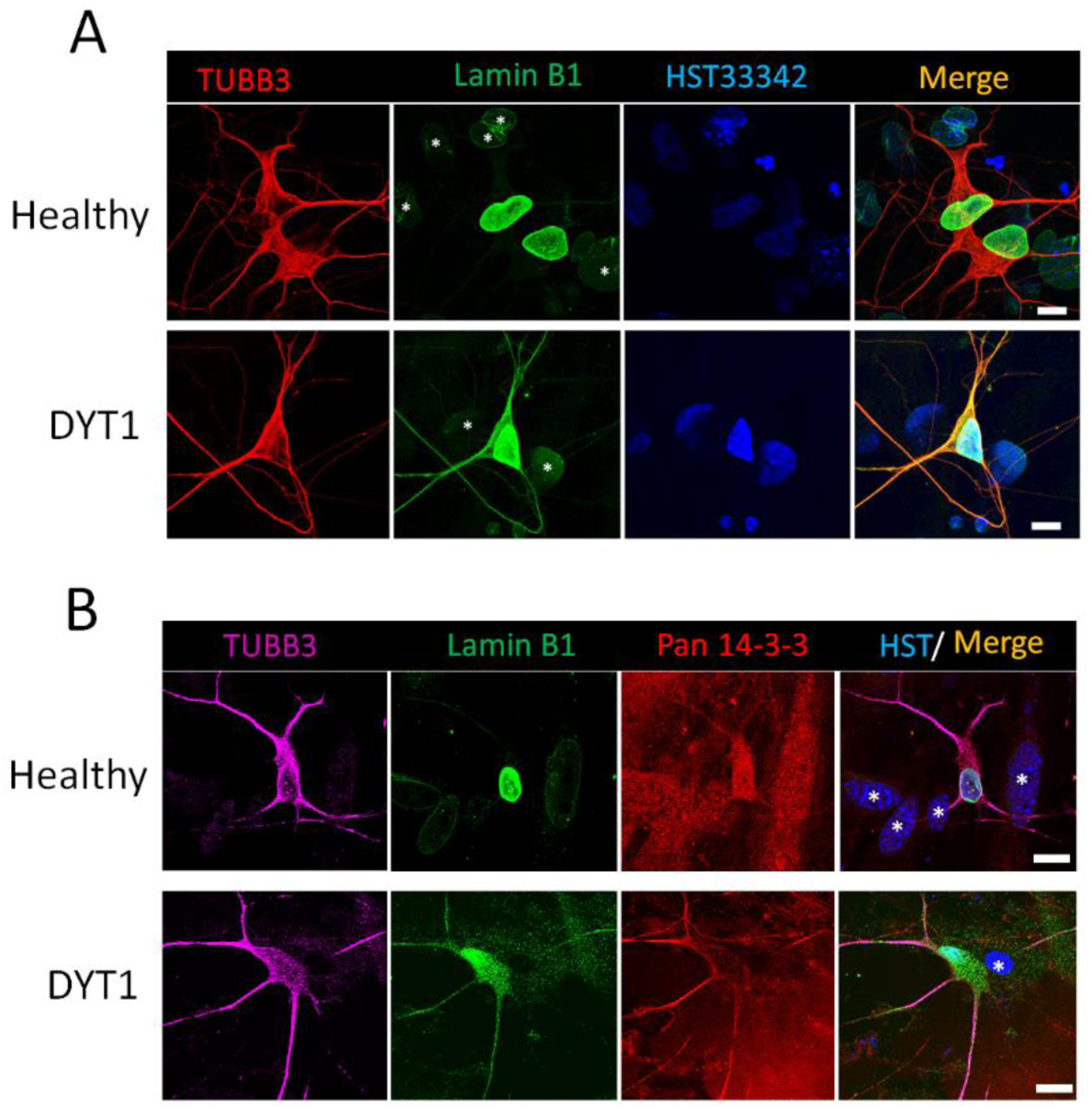
14-3-3 proteins colocalize with cytoplasmically mislocalized Lamin B1 in DYT1 patient-derived motor neurons. **(A)** Confocal micrographs of motor neurons directly reprogrammed from healthy control (GM00024) and DYT1 patient (GM03211) fibroblasts at 5 weeks post-viral infection (WPI). Asterisks indicate co-cultured astrocytes. Scale bars: 20 µm. **(B)** Confocal micrographs showing co-staining of Lamin B1 and pan-14-3-3 proteins in reprogrammed neurons. Asterisks indicate co-cultured astrocytes. The pan-14-3-3 antibody, which recognizes all isotypes, stains both neurons and astrocytes, resulting in high background signal. Nevertheless, 14-3-3 proteins are enriched and colocalize with mislocalized cytoplasmic Lamin B1 in neuronal processes. Scale bars: 20 µm.

**Figure S6.**
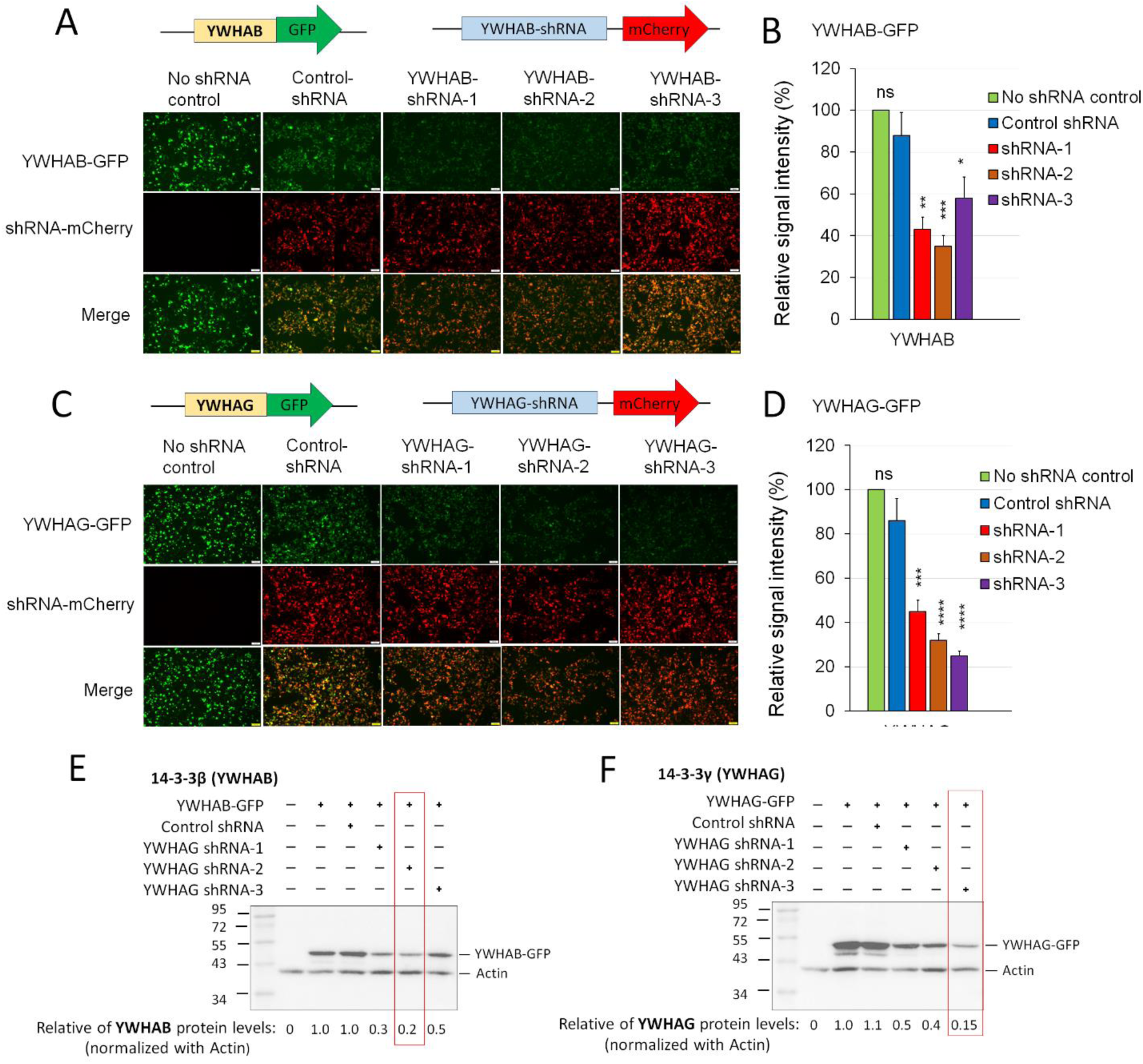
Design and validation of lentiviral vectors to modulate 14-3-3 expression. (**A**) Fluorescence micrographs of HEK cells co-transfected with lentiviral vectors expressing GFP-tagged YWHAB and YWHAB-targeting shRNAs. Three shRNAs target different sites within the coding region of the YWHAB gene. A scrambled sequence was used as a control shRNA. Transfected cells are marked by mCherry, and GFP signal intensity reflects the relative expression level of YWHAB-GFP. Scale bar: 100 µm. (**B**) Quantification of YWHAB-GFP fluorescence intensity from panel (A). Compared to control shRNA: ns, not significant; *p < 0.05; **p < 0.01; ***p < 0.001. One-way ANOVA with Dunnett’s multiple comparison test. (**C**) Fluorescence micrographs of HEK cells co-transfected with GFP-tagged YWHAG and YWHAG-targeting shRNAs. Three shRNAs target different regions of the *YWHAG* coding sequence. Scale bar: 100 µm. (**D**) Quantification of YWHAG-GFP fluorescence intensity from panel (C). Compared to control shRNA: ns, not significant; ***p < 0.001; ****p < 0.0001. One-way ANOVA with Dunnett’s multiple comparison test. (**E**) Western blot showing YWHAB-GFP and β-actin protein levels from whole-cell lysates prepared from panel (A). Blots were sequentially probed with anti-GFP and anti-actin antibodies. Relative YWHAB-GFP protein levels are indicated below. YWHAB-shRNA-2 showed the strongest knockdown efficiency. (**F**) Western blot showing YWHAG-GFP and β-actin protein levels from whole-cell lysates prepared from panel (C). Blots were sequentially probed with anti-GFP and anti-actin antibodies. Relative YWHAG-GFP protein levels are indicated below. YWHAG-shRNA-3 showed the strongest knockdown efficiency.

**Figure S7.**
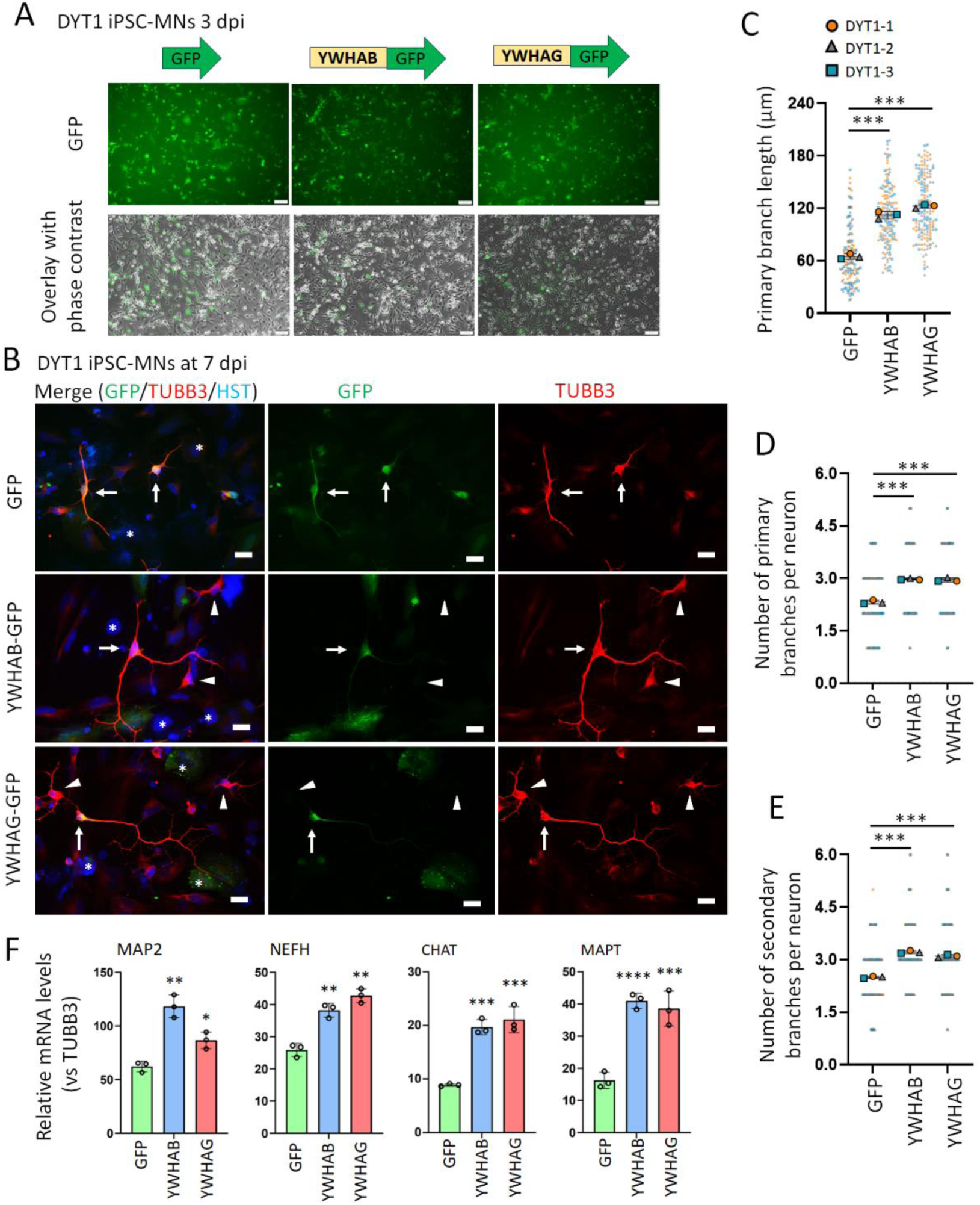
Overexpression of 14-3-3 proteins enhances neurite outgrowth and differentiation in DYT1 neurons. (**A**) Representative micrographs of DYT1 iPSC-MNs (DYT1-1) transduced with lentiviruses expressing GFP, GFP-tagged YWHAB, or YWHAG at 3 dpi. Transduction efficiency and seeded neuron density are comparable across conditions. Scale bar: 100 µm. (**B**) Fluorescence micrographs of DYT1 iPSC-MNs at 7 dpi. Arrows indicate GFP-positive (transduced) neurons; arrowheads indicate GFP-negative (non-transduced) neurons; asterisks denote co-cultured astrocytes. Scale bar: 20 µm. (**C**) Quantification of relative neurite length in DYT1 iPSC-MNs at 7 dpi. N (objects) = 3. n (Neurons) = 50 for each object highlighted with distinct colors. Compared to control: *** p<0.001. One-way ANOVA with Dunnett’s multiple comparison test. (**D**) Quantification of primary neurite branches per neuron at 7 dpi. N (objects) = 3. n (Neurons) = 50 for each object highlighted with distinct colors. Compared to control: *** p<0.001. One-way ANOVA with Dunnett’s multiple comparison test. (**E**) Quantification of secondary neurite branches per neuron at 7 dpi. N (objects) = 3. n (Neurons) = 50 for each object highlighted with distinct colors. Compared to control: *** p<0.001. One-way ANOVA with Dunnett’s multiple comparison test. (**F**) Relative mRNA levels in DYT1-MNs at 7 dpi analyzed by RT-PCR. *n* = 3 biological replicates. Compared to control: * p<0.05, ** p<0.01, *** p<0.001, **** p<0.0001. One-way ANOVA with Dunnett’s multiple comparison test.

**Figure S8.**
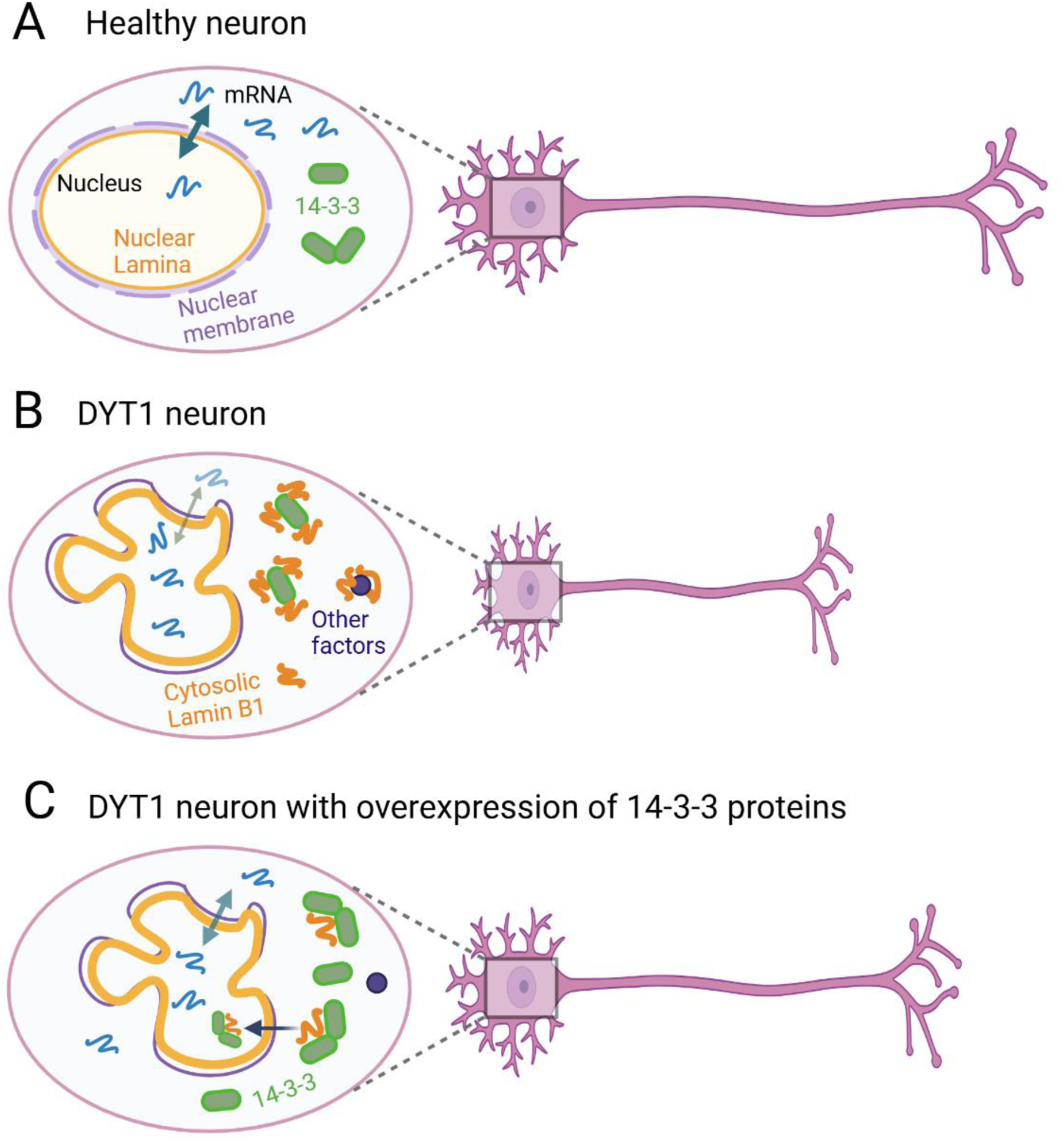
Summary of Lamin B1 dysregulation and its interaction with 14-3-3 proteins in DYT1 neurons. **(A)** A cartoon illustrates a healthy neuron with efficient nucleocytoplasmic transport and normal expression and subcellular localization of nuclear Lamin B1 and 14-3-3 proteins, supporting intrinsic programs that regulate neurite outgrowth, maturation, and neuronal function. **(B)** A cartoon depicts cellular deficits in DYT1 neurons, including nuclear deformation, thickened nuclear lamina, and impaired nucleocytoplasmic transport, resulting in nuclear mRNA accumulation and protein mislocalization. Mislocalized cytoplasmic Lamin B1 strongly binds 14-3-3 proteins, disrupting critical factors and signaling pathways essential for neurodevelopment and neuronal function, ultimately leading to neuronal dysfunction. **(C)** A cartoon shows that overexpression of 14-3-3 proteins rescues DYT1 neuronal deficits by reducing Lamin B1 mislocalization, restoring nucleocytoplasmic transport, and promoting neurite outgrowth and neuronal function.

### Supplementary Tables

**Table S1.**
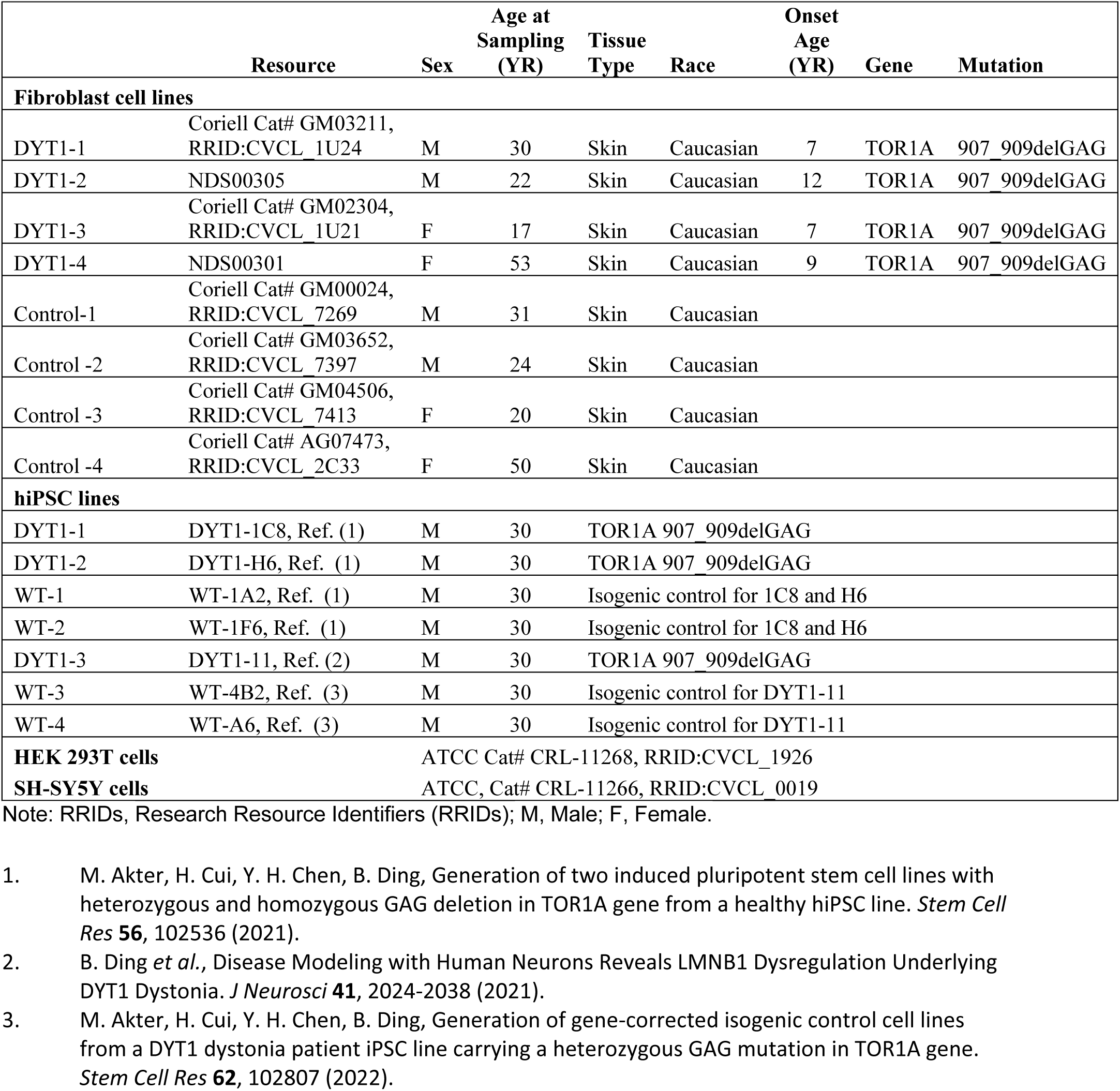
List of cell lines used in this study and their Research Resource Identifiers (RRIDs).

**Table S2.**
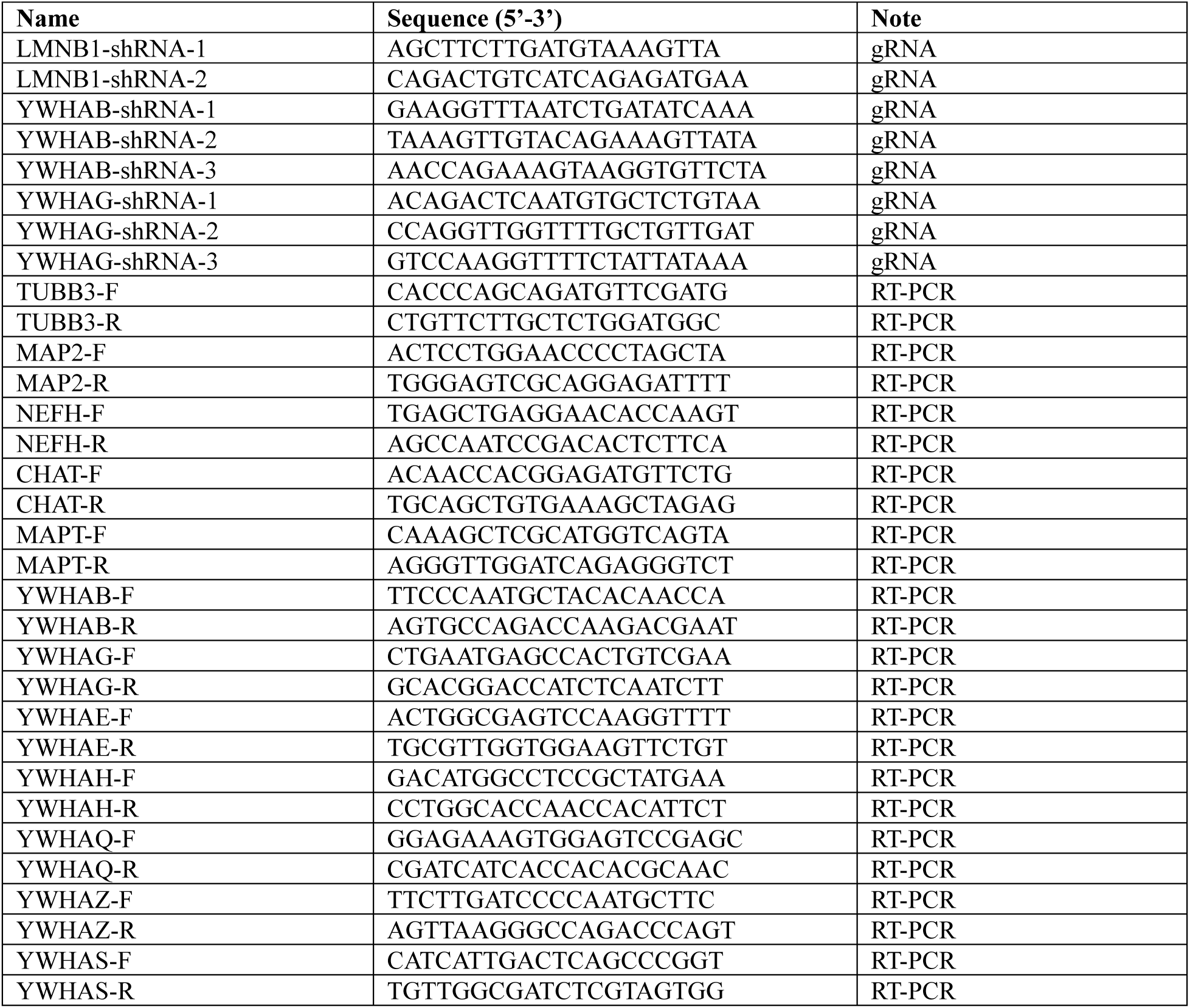
Oligo Sequence.

**Table S3.**
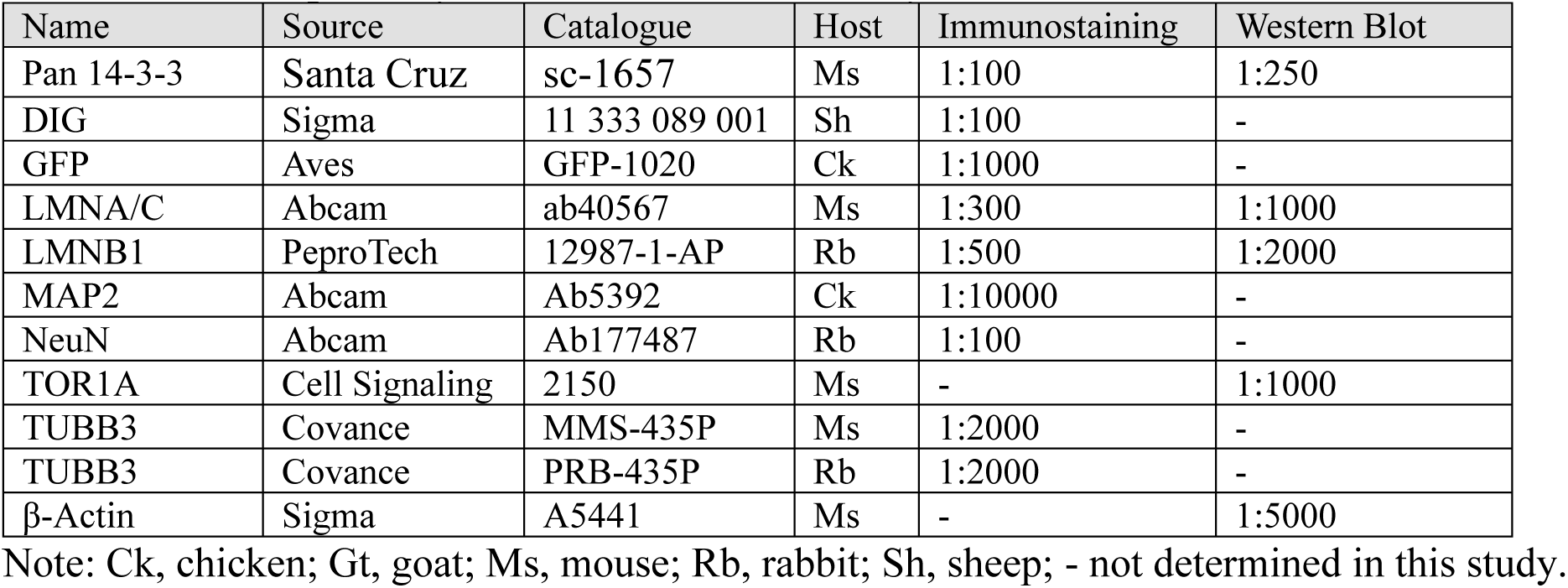
List of primary antibodies used this study.

**Table S4. Co-IP/MS dataset from hiPSC-MNs and the list of reported Lamin B1-interacting proteins.**

**Table S5. GO analysis dataset of Lamin B1-interacting proteins in hiPSC-MNs.**

**Table S6. List of nuclear Lamin B1-associated proteins involved in regulating nuclear transport in iPSC-MNs.**

**Table S7. Categories of neuron-specific proteins associated with nuclear Lamin B1 in iPSC-MNs.**

**Table S8. List of proteins significantly changed the interaction with cytoplasmic Lamin B1 in iPSC-MNs**

**Table S9. List of proteins significantly changed the interaction with cytoplasmic Lamin B1 in SY5Y-Neurons.**

Note: Supplemental tables S4-9 were uploaded separately as excel files.

## Notes

### Competing Interest Statement

The authors have declared no competing interest.

## REFERENCES

1. Turgay, Y., Eibauer, M., Goldman, A.E., Shimi, T., Khayat, M., Ben-Harush, K., Dubrovsky-Gaupp, A., Sapra, K.T., Goldman, R.D., and Medalia, O. (2017). The molecular architecture of lamins in somatic cells. Nature 543, 261–264.

2. Wong, X., Melendez-Perez, A.J., and Reddy, K.L. (2022). The Nuclear Lamina. Cold Spring Harb Perspect Biol 14.

3. Sobo, J.M., Alagna, N.S., Sun, S.X., Wilson, K.L., and Reddy, K.L. (2024). Lamins: The backbone of the nucleocytoskeleton interface. Curr Opin Cell Biol 86, 102313. 10.1016/j.ceb.2023.102313.

4. Yuan, X., Cai, B., Hamamura, Y., Schnittger, A., and Yang, C. (2025). SCF(RMF)-dependent degradation of the nuclear lamina releases the somatic chromatin mobility restriction for meiotic recombination. Sci Adv 11, eadr4567. 10.1126/sciadv.adr4567.

5. Madsen-Osterbye, J., Abdelhalim, M., Pickering, S.H., and Collas, P. (2023). Gene Regulatory Interactions at Lamina-Associated Domains. Genes (Basel) 14. 10.3390/genes14020334.

6. Dechat, T., Pfleghaar, K., Sengupta, K., Shimi, T., Shumaker, D.K., Solimando, L., and Goldman, R.D. (2008). Nuclear lamins: major factors in the structural organization and function of the nucleus and chromatin. Genes Dev 22, 832–853.

7. Capell, B.C., and Collins, F.S. (2006). Human laminopathies: nuclei gone genetically awry. Nat Rev Genet 7, 940–952.

8. Shah, P., Wolf, K., and Lammerding, J. (2017). Bursting the Bubble - Nuclear Envelope Rupture as a Path to Genomic Instability? Trends Cell Biol 27, 546–555.

9. Shin, J.Y., and Worman, H.J. (2022). Molecular Pathology of Laminopathies. Annu Rev Pathol 17, 159–180. 10.1146/annurev-pathol-042220-034240.

10. Lv, T., Wang, C., Zhou, J., Feng, X., Zhang, L., and Fan, Z. (2024). Mechanism and role of nuclear laminin B1 in cell senescence and malignant tumors. Cell Death Discov 10, 269. 10.1038/s41420-024-02045-9.

11. Pennarun, G., Picotto, J., Etourneaud, L., Redavid, A.R., Certain, A., Gauthier, L.R., Fontanilla-Ramirez, P., Busso, D., Chabance-Okumura, C., Theze, B., et al. (2021). Increase in lamin B1 promotes telomere instability by disrupting the shelterin complex in human cells. Nucleic Acids Res 49, 9886–9905. 10.1093/nar/gkab761.

12. Sandoval, A., Garrido, E., Camacho, J., Magana, J.J., and Cisneros, B. (2024). Altered expression and localization of nuclear envelope proteins in a prostate cancer cell system. Mol Biol Rep 51, 898. 10.1007/s11033-024-09836-4.

13. Lange, J., Wood-Kaczmar, A., Ali, A., Farag, S., Ghosh, R., Parker, J., Casey, C., Uno, Y., Kunugi, A., Ferretti, P., et al. (2021). Mislocalization of Nucleocytoplasmic Transport Proteins in Human Huntington’s Disease PSC-Derived Striatal Neurons. Front Cell Neurosci 15, 742763. 10.3389/fncel.2021.742763.

14. Solovei, I., Wang, A.S., Thanisch, K., Schmidt, C.S., Krebs, S., Zwerger, M., Cohen, T.V., Devys, D., Foisner, R., Peichl, L., et al. (2013). LBR and lamin A/C sequentially tether peripheral heterochromatin and inversely regulate differentiation. Cell 152, 584–598. 10.1016/j.cell.2013.01.009.

15. Swift, J., Ivanovska, I.L., Buxboim, A., Harada, T., Dingal, P.C., Pinter, J., Pajerowski, J.D., Spinler, K.R., Shin, J.W., Tewari, M., et al. (2013). Nuclear lamin-A scales with tissue stiffness and enhances matrix-directed differentiation. Science 341, 1240104. 10.1126/science.1240104.

16. Dreesen, O., Chojnowski, A., Ong, P.F., Zhao, T.Y., Common, J.E., Lunny, D., Lane, E.B., Lee, S.J., Vardy, L.A., Stewart, C.L., and Colman, A. (2013). Lamin B1 fluctuations have differential effects on cellular proliferation and senescence. J Cell Biol 200, 605–617.

17. Evangelisti, C., Rusciano, I., Mongiorgi, S., Ramazzotti, G., Lattanzi, G., Manzoli, L., Cocco, L., and Ratti, S. (2022). The wide and growing range of lamin B-related diseases: from laminopathies to cancer. Cell Mol Life Sci 79, 126.

18. Shah, P.P., Donahue, G., Otte, G.L., Capell, B.C., Nelson, D.M., Cao, K., Aggarwala, V., Cruickshanks, H.A., Rai, T.S., McBryan, T., et al. (2013). Lamin B1 depletion in senescent cells triggers large-scale changes in gene expression and the chromatin landscape. Genes Dev 27, 1787–1799.

19. Young, S.G., Jung, H.J., Lee, J.M., and Fong, L.G. (2014). Nuclear lamins and neurobiology. Mol Cell Biol 34, 2776–2785.

20. Butin-Israeli, V., Adam, S.A., Jain, N., Otte, G.L., Neems, D., Wiesmuller, L., Berger, S.L., and Goldman, R.D. (2015). Role of lamin b1 in chromatin instability. Mol Cell Biol 35, 884–898.

21. Kim, Y. (2023). The impact of altered lamin B1 levels on nuclear lamina structure and function in aging and human diseases. Curr Opin Cell Biol 85, 102257. 10.1016/j.ceb.2023.102257.

22. Pappas, S.S., Liang, C.C., Kim, S., Rivera, C.O., and Dauer, W.T. (2018). TorsinA dysfunction causes persistent neuronal nuclear pore defects. Hum Mol Genet 27, 407–420.

23. Dauer, W. (2014). Inherited isolated dystonia: clinical genetics and gene function. Neurotherapeutics 11, 807–816.

24. Li, J., Levin, D.S., Kim, A.J., Pappas, S.S., and Dauer, W.T. (2021). TorsinA restoration in a mouse model identifies a critical therapeutic window for DYT1 dystonia. J Clin Invest.

25. Gonzalez-Alegre, P. (2019). Advances in molecular and cell biology of dystonia: Focus on torsinA. Neurobiol Dis 127, 233–241.

26. Keller Sarmiento, I.J., and Mencacci, N.E. (2021). Genetic Dystonias: Update on Classification and New Genetic Discoveries. Curr Neurol Neurosci Rep 21, 8.

27. Ozelius, L.J., Hewett, J.W., Page, C.E., Bressman, S.B., Kramer, P.L., Shalish, C., de Leon, D., Brin, M.F., Raymond, D., Corey, D.P., et al. (1997). The early-onset torsion dystonia gene (DYT1) encodes an ATP-binding protein. Nat Genet 17, 40–48.

28. Charlesworth, G., Bhatia, K.P., and Wood, N.W. (2013). The genetics of dystonia: new twists in an old tale. Brain 136, 2017–2037.

29. Hanson, P.I., and Whiteheart, S.W. (2005). AAA+ proteins: have engine, will work. Nat Rev Mol Cell Biol 6, 519–529.

30. Rampello, A.J., Prophet, S.M., and Schlieker, C. (2020). The Role of Torsin AAA+ Proteins in Preserving Nuclear Envelope Integrity and Safeguarding Against Disease. Biomolecules 10.

31. Ogura, T., and Wilkinson, A.J. (2001). AAA+ superfamily ATPases: common structure--diverse function. Genes Cells 6, 575–597.

32. White, S.R., and Lauring, B. (2007). AAA+ ATPases: achieving diversity of function with conserved machinery. Traffic 8, 1657–1667.

33. Kustedjo, K., Bracey, M.H., and Cravatt, B.F. (2000). Torsin A and its torsion dystonia-associated mutant forms are lumenal glycoproteins that exhibit distinct subcellular localizations. J Biol Chem 275, 27933–27939. 10.1074/jbc.M910025199.

34. Vander Heyden, A.B., Naismith, T.V., Snapp, E.L., and Hanson, P.I. (2011). Static retention of the lumenal monotopic membrane protein torsinA in the endoplasmic reticulum. EMBO J 30, 3217–3231.

35. Naismith, T.V., Heuser, J.E., Breakefield, X.O., and Hanson, P.I. (2004). TorsinA in the nuclear envelope. Proc Natl Acad Sci U S A 101, 7612–7617.

36. Hewett, J.W., Tannous, B., Niland, B.P., Nery, F.C., Zeng, J., Li, Y., and Breakefield, X.O. (2007). Mutant torsinA interferes with protein processing through the secretory pathway in DYT1 dystonia cells. Proc Natl Acad Sci U S A 104, 7271–7276.

37. Hewett, J.W., Nery, F.C., Niland, B., Ge, P., Tan, P., Hadwiger, P., Tannous, B.A., Sah, D.W., and Breakefield, X.O. (2008). siRNA knock-down of mutant torsinA restores processing through secretory pathway in DYT1 dystonia cells. Hum Mol Genet 17, 1436–1445.

38. Beauvais, G., Bode, N.M., Watson, J.L., Wen, H., Glenn, K.A., Kawano, H., Harata, N.C., Ehrlich, M.E., and Gonzalez-Alegre, P. (2016). Disruption of Protein Processing in the Endoplasmic Reticulum of DYT1 Knock-in Mice Implicates Novel Pathways in Dystonia Pathogenesis. J Neurosci 36, 10245–10256.

39. Nery, F.C., Armata, I.A., Farley, J.E., Cho, J.A., Yaqub, U., Chen, P., da Hora, C.C., Wang, Q., Tagaya, M., Klein, C., et al. (2011). TorsinA participates in endoplasmic reticulum-associated degradation. Nat Commun 2, 393.

40. Burdette, A.J., Churchill, P.F., Caldwell, G.A., and Caldwell, K.A. (2010). The early-onset torsion dystonia-associated protein, torsinA, displays molecular chaperone activity in vitro. Cell Stress Chaperones 15, 605–617. 10.1007/s12192-010-0173-2.

41. Chen, P., Burdette, A.J., Porter, J.C., Ricketts, J.C., Fox, S.A., Nery, F.C., Hewett, J.W., Berkowitz, L.A., Breakefield, X.O., Caldwell, K.A., and Caldwell, G.A. (2010). The early-onset torsion dystonia-associated protein, torsinA, is a homeostatic regulator of endoplasmic reticulum stress response. Hum Mol Genet 19, 3502–3515.

42. Nery, F.C., Zeng, J., Niland, B.P., Hewett, J., Farley, J., Irimia, D., Li, Y., Wiche, G., Sonnenberg, A., and Breakefield, X.O. (2008). TorsinA binds the KASH domain of nesprins and participates in linkage between nuclear envelope and cytoskeleton. J Cell Sci 121, 3476–3486.

43. Laudermilch, E., Tsai, P.L., Graham, M., Turner, E., Zhao, C., and Schlieker, C. (2016). Dissecting Torsin/cofactor function at the nuclear envelope: a genetic study. Mol Biol Cell 27, 3964–3971.

44. Ding, B., Tang, Y., Ma, S., Akter, M., Liu, M.L., Zang, T., and Zhang, C.L. (2021). Disease Modeling with Human Neurons Reveals LMNB1 Dysregulation Underlying DYT1 Dystonia. J Neurosci 41, 2024–2038.

45. Prophet, S.M., Rampello, A.J., Niescier, R.F., Gentile, J.E., Mallik, S., Koleske, A.J., and Schlieker, C. (2022). Atypical nuclear envelope condensates linked to neurological disorders reveal nucleoporin-directed chaperone activities. Nat Cell Biol.

46. Rampello, A.J., Laudermilch, E., Vishnoi, N., Prophet, S.M., Shao, L., Zhao, C., Lusk, C.P., and Schlieker, C. (2020). Torsin ATPase deficiency leads to defects in nuclear pore biogenesis and sequestration of MLF2. J Cell Biol 219.

47. VanGompel, M.J., Nguyen, K.C., Hall, D.H., Dauer, W.T., and Rose, L.S. (2015). A novel function for the Caenorhabditis elegans torsin OOC-5 in nucleoporin localization and nuclear import. Mol Biol Cell 26, 1752–1763.

48. Kim, S., Phan, S., Tran, H.T., Shaw, T.R., Shahmoradian, S.H., Ellisman, M.H., Veatch, S.L., Barmada, S.J., Pappas, S.S., and Dauer, W.T. (2024). TorsinA is essential for neuronal nuclear pore complex localization and maturation. Nat Cell Biol 26, 1482–1495. 10.1038/s41556-024-01480-1.

49. Jokhi, V., Ashley, J., Nunnari, J., Noma, A., Ito, N., Wakabayashi-Ito, N., Moore, M.J., and Budnik, V. (2013). Torsin mediates primary envelopment of large ribonucleoprotein granules at the nuclear envelope. Cell Rep 3, 988–995.

50. Maric, M., Shao, J., Ryan, R.J., Wong, C.S., Gonzalez-Alegre, P., and Roller, R.J. (2011). A functional role for TorsinA in herpes simplex virus 1 nuclear egress. J Virol 85, 9667–9679.

51. Pocratsky, A.M., Nascimento, F., Ozyurt, M.G., White, I.J., Sullivan, R., O’Callaghan, B.J., Smith, C.C., Surana, S., Beato, M., and Brownstone, R.M. (2023). Pathophysiology of Dyt1-Tor1a dystonia in mice is mediated by spinal neural circuit dysfunction. Sci Transl Med 15, eadg3904.

52. Torres, G.E., Sweeney, A.L., Beaulieu, J.M., Shashidharan, P., and Caron, M.G. (2004). Effect of torsinA on membrane proteins reveals a loss of function and a dominant-negative phenotype of the dystonia-associated DeltaE-torsinA mutant. Proc Natl Acad Sci U S A 101, 15650–15655.

53. Goodchild, R.E., Kim, C.E., and Dauer, W.T. (2005). Loss of the dystonia-associated protein torsinA selectively disrupts the neuronal nuclear envelope. Neuron 48, 923–932.

54. Zhao, C., Brown, R.S., Chase, A.R., Eisele, M.R., and Schlieker, C. (2013). Regulation of Torsin ATPases by LAP1 and LULL1. Proc Natl Acad Sci U S A 110, E1545–1554.

55. Tanabe, L.M., Liang, C.C., and Dauer, W.T. (2016). Neuronal Nuclear Membrane Budding Occurs during a Developmental Window Modulated by Torsin Paralogs. Cell Rep 16, 3322–3333. 10.1016/j.celrep.2016.08.044.

56. Prophet, S.M., Rampello, A.J., Niescier, R.F., Gentile, J.E., Mallik, S., Koleske, A.J., and Schlieker, C. (2022). Atypical nuclear envelope condensates linked to neurological disorders reveal nucleoporin-directed chaperone activities. Nat Cell Biol 24, 1630–1641. 10.1038/s41556-022-01001-y.

57. Ding, B. (2021). Generation of patient-specific motor neurons in modeling movement diseases. Neural Regen Res 16, 1799–1800.

58. Sepehrimanesh, M., Akter, M., and Ding, B. (2021). Direct conversion of adult fibroblasts into motor neurons. STAR Protoc 2, 100917.

59. Akter, M., Cui, H., Sepehrimanesh, M., Hosain, M.A., and Ding, B. (2022). Generation of highly pure motor neurons from human induced pluripotent stem cells. STAR Protoc 3, 101223.

60. Akter, M., Cui, H., Chen, Y.H., and Ding, B. (2021). Generation of two induced pluripotent stem cell lines with heterozygous and homozygous GAG deletion in TOR1A gene from a healthy hiPSC line. Stem Cell Res 56, 102536.

61. Akter, M., Cui, H., Chen, Y.H., and Ding, B. (2022). Generation of gene-corrected isogenic control cell lines from a DYT1 dystonia patient iPSC line carrying a heterozygous GAG mutation in TOR1A gene. Stem Cell Res 62, 102807. 10.1016/j.scr.2022.102807.

62. Akter, M., Cui, H., Hosain, M.A., Liu, J., Duan, Y., and Ding, B. (2024). RANBP17 Overexpression Restores Nucleocytoplasmic Transport and Ameliorates Neurodevelopment in Induced DYT1 Dystonia Motor Neurons. J Neurosci 44. 10.1523/JNEUROSCI.1728-23.2024.

63. Ding, B. (2022). Novel insights into the pathogenesis of DYT1 dystonia from induced patient-derived neurons. Neural Regen Res 17, 561–562.

64. Cho, E., and Park, J.Y. (2020). Emerging roles of 14-3-3gamma in the brain disorder. BMB Rep 53, 500–511. 10.5483/BMBRep.2020.53.10.158.

65. Cornell, B., and Toyo-Oka, K. (2017). 14-3-3 Proteins in Brain Development: Neurogenesis, Neuronal Migration and Neuromorphogenesis. Front Mol Neurosci 10, 318.

66. Levy, D.L., and Heald, R. (2010). Nuclear size is regulated by importin alpha and Ntf2 in Xenopus. Cell 143, 288–298. 10.1016/j.cell.2010.09.012.

67. Vukovic, L.D., Jevtic, P., Zhang, Z., Stohr, B.A., and Levy, D.L. (2016). Nuclear size is sensitive to NTF2 protein levels in a manner dependent on Ran binding. J Cell Sci 129, 1115–1127. 10.1242/jcs.181263.

68. Cui H., S.M., Coutee C.,, Akter M., Hosain A., and B., D. (2022). Protocol to image and quantify nucleocytoplasmic transport in cultured cells using fluorescent in situ hybridization and a dual reporter sytem. STAR Protoc 3, 101813.

69. Mertens, J., Paquola, A.C., Ku, M., Hatch, E., Bohnke, L., Ladjevardi, S., McGrath, S., Campbell, B., Lee, H., Herdy, J.R., et al. (2015). Directly Reprogrammed Human Neurons Retain Aging-Associated Transcriptomic Signatures and Reveal Age-Related Nucleocytoplasmic Defects. Cell Stem Cell 17, 705–718.

70. Ding, B., Akter, M., and Zhang, C.L. (2020). Differential Influence of Sample Sex and Neuronal Maturation on mRNA and Protein Transport in Induced Human Neurons. Front Mol Neurosci 13, 46.

71. Akter, M., Sepehrimanesh, M., Xu, W., and Ding, B. (2024). Assembling a Coculture System to Prepare Highly Pure Induced Pluripotent Stem Cell-Derived Neurons at Late Maturation Stages. eNeuro 11. 10.1523/ENEURO.0165-24.2024.

72. Sepehrimanesh, M., and Ding, B. (2020). Generation and optimization of highly pure motor neurons from human induced pluripotent stem cells via lentiviral delivery of transcription factors. Am J Physiol Cell Physiol 319, C771–C780.

73. Ahn, J., Lee, J., Jeong, S., Kang, S.M., Park, B.J., and Ha, N.C. (2021). Beta-strand-mediated dimeric formation of the Ig-like domains of human lamin A/C and B1. Biochem Biophys Res Commun 550, 191–196. 10.1016/j.bbrc.2021.02.102.

74. Ruan, J., Xu, C., Bian, C., Lam, R., Wang, J.P., Kania, J., Min, J., and Zang, J. (2012). Crystal structures of the coil 2B fragment and the globular tail domain of human lamin B1. FEBS Lett 586, 314–318. 10.1016/j.febslet.2012.01.007.

75. Jumper, J., Evans, R., Pritzel, A., Green, T., Figurnov, M., Ronneberger, O., Tunyasuvunakool, K., Bates, R., Zidek, A., Potapenko, A., et al. (2021). Highly accurate protein structure prediction with AlphaFold. Nature 596, 583–589. 10.1038/s41586-021-03819-2.

76. Abramson, J., Adler, J., Dunger, J., Evans, R., Green, T., Pritzel, A., Ronneberger, O., Willmore, L., Ballard, A.J., Bambrick, J., et al. (2024). Accurate structure prediction of biomolecular interactions with AlphaFold 3. Nature 630, 493–500. 10.1038/s41586-024-07487-w.

77. Akter, M., and Ding, B. (2022). Modeling Movement Disorders via Generation of hiPSC-Derived Motor Neurons. Cells 11.

78. Oughtred, R., Stark, C., Breitkreutz, B.J., Rust, J., Boucher, L., Chang, C., Kolas, N., O’Donnell, L., Leung, G., McAdam, R., et al. (2019). The BioGRID interaction database: 2019 update. Nucleic Acids Res 47, D529–D541. 10.1093/nar/gky1079.

79. Wong, X., Cutler, J.A., Hoskins, V.E., Gordon, M., Madugundu, A.K., Pandey, A., and Reddy, K.L. (2021). Mapping the micro-proteome of the nuclear lamina and lamina-associated domains. Life Sci Alliance 4. 10.26508/lsa.202000774.

80. Cho, N.H., Cheveralls, K.C., Brunner, A.D., Kim, K., Michaelis, A.C., Raghavan, P., Kobayashi, H., Savy, L., Li, J.Y., Canaj, H., et al. (2022). OpenCell: Endogenous tagging for the cartography of human cellular organization. Science 375, eabi6983. 10.1126/science.abi6983.

81. Go, C.D., Knight, J.D.R., Rajasekharan, A., Rathod, B., Hesketh, G.G., Abe, K.T., Youn, J.Y., Samavarchi-Tehrani, P., Zhang, H., Zhu, L.Y., et al. (2021). A proximity-dependent biotinylation map of a human cell. Nature 595, 120–124. 10.1038/s41586-021-03592-2.

82. Krastev, D.B., Li, S., Sun, Y., Wicks, A.J., Hoslett, G., Weekes, D., Badder, L.M., Knight, E.G., Marlow, R., Pardo, M.C., et al. (2022). The ubiquitin-dependent ATPase p97 removes cytotoxic trapped PARP1 from chromatin. Nat Cell Biol 24, 62–73. 10.1038/s41556-021-00807-6.

83. Havugimana, P.C., Goel, R.K., Phanse, S., Youssef, A., Padhorny, D., Kotelnikov, S., Kozakov, D., and Emili, A. (2022). Scalable multiplex co-fractionation/mass spectrometry platform for accelerated protein interactome discovery. Nat Commun 13, 4043. 10.1038/s41467-022-31809-z.

84. Wing, C.E., Fung, H.Y.J., and Chook, Y.M. (2022). Karyopherin-mediated nucleocytoplasmic transport. Nat Rev Mol Cell Biol 23, 307–328. 10.1038/s41580-021-00446-7.

85. Ding, B., and Sepehrimanesh, M. (2021). Nucleocytoplasmic Transport: Regulatory Mechanisms and the Implications in Neurodegeneration. Int J Mol Sci 22.

86. Kadowaki, T., Goldfarb, D., Spitz, L.M., Tartakoff, A.M., and Ohno, M. (1993). Regulation of RNA processing and transport by a nuclear guanine nucleotide release protein and members of the Ras superfamily. EMBO J 12, 2929–2937. 10.1002/j.1460-2075.1993.tb05955.x.

87. Bischoff, F.R., Klebe, C., Kretschmer, J., Wittinghofer, A., and Ponstingl, H. (1994). RanGAP1 induces GTPase activity of nuclear Ras-related Ran. Proc Natl Acad Sci U S A 91, 2587–2591. 10.1073/pnas.91.7.2587.

88. Bischoff, F.R., Krebber, H., Kempf, T., Hermes, I., and Ponstingl, H. (1995). Human RanGTPase-activating protein RanGAP1 is a homologue of yeast Rna1p involved in mRNA processing and transport. Proc Natl Acad Sci U S A 92, 1749–1753. 10.1073/pnas.92.5.1749.

89. Falini, B., Sorcini, D., Perriello, V.M., and Sportoletti, P. (2025). Functions of the native NPM1 protein and its leukemic mutant. Leukemia 39, 276–290. 10.1038/s41375-024-02476-4.

90. Percipalle, P., Raju, C.S., and Fukuda, N. (2009). Actin-associated hnRNP proteins as transacting factors in the control of mRNA transport and localization. RNA Biol 6, 171–174. 10.4161/rna.6.2.8195.

91. He, Y., and Smith, R. (2009). Nuclear functions of heterogeneous nuclear ribonucleoproteins A/B. Cell Mol Life Sci 66, 1239–1256. 10.1007/s00018-008-8532-1.

92. Bassler, J., and Hurt, E. (2019). Eukaryotic Ribosome Assembly. Annu Rev Biochem 88, 281–306. 10.1146/annurev-biochem-013118-110817.

93. Nerurkar, P., Altvater, M., Gerhardy, S., Schutz, S., Fischer, U., Weirich, C., and Panse, V.G. (2015). Eukaryotic Ribosome Assembly and Nuclear Export. Int Rev Cell Mol Biol 319, 107–140. 10.1016/bs.ircmb.2015.07.002.

94. Gigante, C.M., Dibattista, M., Dong, F.N., Zheng, X., Yue, S., Young, S.G., Reisert, J., Zheng, Y., and Zhao, H. (2017). Lamin B1 is required for mature neuron-specific gene expression during olfactory sensory neuron differentiation. Nat Commun 8, 15098. 10.1038/ncomms15098.

95. Chen, N.Y., Yang, Y., Weston, T.A., Belling, J.N., Heizer, P., Tu, Y., Kim, P., Edillo, L., Jonas, S.J., Weiss, P.S., et al. (2019). An absence of lamin B1 in migrating neurons causes nuclear membrane ruptures and cell death. Proc Natl Acad Sci U S A 116, 25870–25879. 10.1073/pnas.1917225116.

96. Koufi, F.D., Neri, I., Ramazzotti, G., Rusciano, I., Mongiorgi, S., Marvi, M.V., Fazio, A., Shin, M., Kosodo, Y., Cani, I., et al. (2023). Lamin B1 as a key modulator of the developing and aging brain. Front Cell Neurosci 17, 1263310. 10.3389/fncel.2023.1263310.

97. Giacomini, C., Mahajani, S., Ruffilli, R., Marotta, R., and Gasparini, L. (2016). Lamin B1 protein is required for dendrite development in primary mouse cortical neurons. Mol Biol Cell 27, 35–47. 10.1091/mbc.E15-05-0307.

98. Nolan, M., Talbot, K., and Ansorge, O. (2016). Pathogenesis of FUS-associated ALS and FTD: insights from rodent models. Acta Neuropathol Commun 4, 99. 10.1186/s40478-016-0358-8.

99. Kwiatkowski, T.J., Jr., Bosco, D.A., Leclerc, A.L., Tamrazian, E., Vanderburg, C.R., Russ, C., Davis, A., Gilchrist, J., Kasarskis, E.J., Munsat, T., et al. (2009). Mutations in the FUS/TLS gene on chromosome 16 cause familial amyotrophic lateral sclerosis. Science 323, 1205–1208. 10.1126/science.1166066.

100. Vance, C., Rogelj, B., Hortobagyi, T., De Vos, K.J., Nishimura, A.L., Sreedharan, J., Hu, X., Smith, B., Ruddy, D., Wright, P., et al. (2009). Mutations in FUS, an RNA processing protein, cause familial amyotrophic lateral sclerosis type 6. Science 323, 1208–1211. 10.1126/science.1165942.

101. Lim, Y.W., James, D., Huang, J., and Lee, M. (2020). The Emerging Role of the RNA-Binding Protein SFPQ in Neuronal Function and Neurodegeneration. Int J Mol Sci 21. 10.3390/ijms21197151.

102. Khodyreva, S.N., Dyrkheeva, N.S., and Lavrik, O.I. (2024). Proteins Associated with Neurodegenerative Diseases: Link to DNA Repair. Biomedicines 12. 10.3390/biomedicines12122808.

103. Dong, Y.T., Luo, X., Zhang, L.L., Gong, Y.M., Wang, D., Zhong, D.L., Li, Y.X., Ma, X.M., Jin, R.J., and Li, J. (2025). Genetic colocalization of cathepsins H, D, and L1 with Alzheimer’s disease: Implications for biomarker and therapeutic target discovery. J Alzheimers Dis 104, 61–72. 10.1177/13872877251314058.

104. Patzke, C., Han, Y., Covy, J., Yi, F., Maxeiner, S., Wernig, M., and Sudhof, T.C. (2015). Analysis of conditional heterozygous STXBP1 mutations in human neurons. J Clin Invest 125, 3560–3571. 10.1172/JCI78612.

105. O’Brien, S., Ng-Cordell, E., Study, D.D.D., Astle, D.E., Scerif, G., and Baker, K. (2019). STXBP1-associated neurodevelopmental disorder: a comparative study of behavioural characteristics. J Neurodev Disord 11, 17. 10.1186/s11689-019-9278-9.

106. De Paola, E., Forcina, L., Pelosi, L., Pisu, S., La Rosa, P., Cesari, E., Nicoletti, C., Madaro, L., Mercatelli, N., Biamonte, F., et al. (2020). Sam68 splicing regulation contributes to motor unit establishment in the postnatal skeletal muscle. Life Sci Alliance 3. 10.26508/lsa.201900637.

107. Farini, D., Cesari, E., Weatheritt, R.J., La Sala, G., Naro, C., Pagliarini, V., Bonvissuto, D., Medici, V., Guerra, M., Di Pietro, C., et al. (2020). A Dynamic Splicing Program Ensures Proper Synaptic Connections in the Developing Cerebellum. Cell Rep 31, 107703. 10.1016/j.celrep.2020.107703.

108. Witte, H., Schreiner, D., and Scheiffele, P. (2019). A Sam68-dependent alternative splicing program shapes postsynaptic protein complexes. Eur J Neurosci 49, 1436–1453. 10.1111/ejn.14332.

109. Carecchio, M., Zorzi, G., Ragona, F., Zibordi, F., and Nardocci, N. (2018). ATP1A3-related disorders: An update. Eur J Paediatr Neurol 22, 257–263. 10.1016/j.ejpn.2017.12.009.

110. Roubergue, A., Roze, E., Vuillaumier-Barrot, S., Fontenille, M.J., Meneret, A., Vidailhet, M., Fontaine, B., Doummar, D., Philibert, B., Riant, F., and Nicole, S. (2013). The multiple faces of the ATP1A3-related dystonic movement disorder. Mov Disord 28, 1457–1459. 10.1002/mds.25396.

111. Bell, M., and Zempel, H. (2022). SH-SY5Y-derived neurons: a human neuronal model system for investigating TAU sorting and neuronal subtype-specific TAU vulnerability. Rev Neurosci 33, 1–15.

112. Kovalevich, J., and Langford, D. (2013). Considerations for the use of SH-SY5Y neuroblastoma cells in neurobiology. Methods Mol Biol 1078, 9–21.

113. Aitken, A. (2006). 14-3-3 proteins: a historic overview. Semin Cancer Biol 16, 162–172.

114. Fan, X., Cui, L., Zeng, Y., Song, W., Gaur, U., and Yang, M. (2019). 14-3-3 Proteins Are on the Crossroads of Cancer, Aging, and Age-Related Neurodegenerative Disease. Int J Mol Sci 20. 10.3390/ijms20143518.

115. Roberts, B.J., Reddy, R., and Wahl, J.K., 3rd (2013). Stratifin (14-3-3 sigma) limits plakophilin-3 exchange with the desmosomal plaque. PLoS One 8, e77012. 10.1371/journal.pone.0077012.

116. Tamburrino, A., and Decressac, M. (2016). Aged and Diseased Neurons Get Lost in Transport. Trends Neurosci 39, 199–201. 10.1016/j.tins.2016.02.007.

117. Nmezi, B., Xu, J., Fu, R., Armiger, T.J., Rodriguez-Bey, G., Powell, J.S., Ma, H., Sullivan, M., Tu, Y., Chen, N.Y., et al. (2019). Concentric organization of A- and B-type lamins predicts their distinct roles in the spatial organization and stability of the nuclear lamina. Proc Natl Acad Sci U S A 116, 4307–4315.

118. Goodchild, R.E., and Dauer, W.T. (2004). Mislocalization to the nuclear envelope: an effect of the dystonia-causing torsinA mutation. Proc Natl Acad Sci U S A 101, 847–852.

119. Kaneshiro, J.M., Capitanio, J.S., and Hetzer, M.W. (2023). Lamin B1 overexpression alters chromatin organization and gene expression. Nucleus 14, 2202548. 10.1080/19491034.2023.2202548.

120. Vergnes, L., Peterfy, M., Bergo, M.O., Young, S.G., and Reue, K. (2004). Lamin B1 is required for mouse development and nuclear integrity. Proc Natl Acad Sci U S A 101, 10428–10433.

121. Coffinier, C., Jung, H.J., Nobumori, C., Chang, S., Tu, Y., Barnes, R.H., 2nd, Yoshinaga, Y., de Jong, P.J., Vergnes, L., Reue, K., et al. (2011). Deficiencies in lamin B1 and lamin B2 cause neurodevelopmental defects and distinct nuclear shape abnormalities in neurons. Mol Biol Cell 22, 4683–4693.

122. Garcia-Forn, M., Castany-Pladevall, C., Golbano, A., Perez-Perez, J., Brito, V., Kulisevsky, J., and Perez-Navarro, E. (2023). Lamin B1 and nuclear morphology in peripheral cells as new potential biomarkers to follow treatment response in Huntington’s disease. Clin Transl Med 13, e1154. 10.1002/ctm2.1154.

123. Chi, Y.H., Wang, W.P., Hung, M.C., Liou, G.G., Wang, J.Y., and Chao, P.G. (2022). Deformation of the nucleus by TGFbeta1 via the remodeling of nuclear envelope and histone isoforms. Epigenetics Chromatin 15, 1.

124. Neelam, S., Richardson, B., Barker, R., Udave, C., Gilroy, S., Cameron, M.J., Levine, H.G., and Zhang, Y. (2020). Changes in Nuclear Shape and Gene Expression in Response to Simulated Microgravity Are LINC Complex-Dependent. Int J Mol Sci 21.

125. Foote, M., and Zhou, Y. (2012). 14-3-3 proteins in neurological disorders. Int J Biochem Mol Biol 3, 152–164.

126. Oh, H.S., Urey, D.Y., Karlsson, L., Zhu, Z., Shen, Y., Farinas, A., Timsina, J., Duggan, M.R., Chen, J., Guldner, I.H., et al. (2025). A cerebrospinal fluid synaptic protein biomarker for prediction of cognitive resilience versus decline in Alzheimer’s disease. Nat Med 31, 1592–1603. 10.1038/s41591-025-03565-2.

127. Sluchanko, N.N., and Gusev, N.B. (2017). Moonlighting chaperone-like activity of the universal regulatory 14-3-3 proteins. FEBS J 284, 1279–1295. 10.1111/febs.13986.

128. Segal, D., Maier, S., Mastromarco, G.J., Qian, W.W., Nabeel-Shah, S., Lee, H., Moore, G., Lacoste, J., Larsen, B., Lin, Z.Y., et al. (2023). A central chaperone-like role for 14-3-3 proteins in human cells. Mol Cell 83, 974–993 e915. 10.1016/j.molcel.2023.02.018.

129. Huang, X., Zheng, Z., Wu, Y., Gao, M., Su, Z., and Huang, Y. (2022). 14-3-3 Proteins are Potential Regulators of Liquid-Liquid Phase Separation. Cell Biochem Biophys 80, 277–293. 10.1007/s12013-022-01067-3.

130. Muslin, A.J., and Xing, H. (2000). 14-3-3 proteins: regulation of subcellular localization by molecular interference. Cell Signal 12, 703–709. 10.1016/s0898-6568(00)00131-5.

131. Obsil, T., Ghirlando, R., Klein, D.C., Ganguly, S., and Dyda, F. (2001). Crystal structure of the 14-3-3zeta:serotonin N-acetyltransferase complex. a role for scaffolding in enzyme regulation. Cell 105, 257–267. 10.1016/s0092-8674(01)00316-6.

132. Reincke, M., Sbiera, S., Hayakawa, A., Theodoropoulou, M., Osswald, A., Beuschlein, F., Meitinger, T., Mizuno-Yamasaki, E., Kawaguchi, K., Saeki, Y., et al. (2015). Mutations in the deubiquitinase gene USP8 cause Cushing’s disease. Nat Genet 47, 31–38. 10.1038/ng.3166.

133. Park, E., Rawson, S., Li, K., Kim, B.W., Ficarro, S.B., Pino, G.G., Sharif, H., Marto, J.A., Jeon, H., and Eck, M.J. (2019). Architecture of autoinhibited and active BRAF-MEK1-14-3-3 complexes. Nature 575, 545–550. 10.1038/s41586-019-1660-y.

134. Grozinger, C.M., and Schreiber, S.L. (2000). Regulation of histone deacetylase 4 and 5 and transcriptional activity by 14-3-3-dependent cellular localization. Proc Natl Acad Sci U S A 97, 7835–7840. 10.1073/pnas.140199597.

135. Fu, H., Subramanian, R.R., and Masters, S.C. (2000). 14-3-3 proteins: structure, function, and regulation. Annu Rev Pharmacol Toxicol 40, 617–647. 10.1146/annurev.pharmtox.40.1.617.

136. Zhang, J., Weinrich, J.A.P., Russ, J.B., Comer, J.D., Bommareddy, P.K., DiCasoli, R.J., Wright, C.V.E., Li, Y., van Roessel, P.J., and Kaltschmidt, J.A. (2017). A Role for Dystonia-Associated Genes in Spinal GABAergic Interneuron Circuitry. Cell Rep 21, 666–678.

137. Downs, A.M., Roman, K.M., Campbell, S.A., Pisani, A., Hess, E.J., and Bonsi, P. (2019). The neurobiological basis for novel experimental therapeutics in dystonia. Neurobiol Dis 130, 104526.

138. Maltese, M., Stanic, J., Tassone, A., Sciamanna, G., Ponterio, G., Vanni, V., Martella, G., Imbriani, P., Bonsi, P., Mercuri, N.B., et al. (2018). Early structural and functional plasticity alterations in a susceptibility period of DYT1 dystonia mouse striatum. Elife 7.

139. Ip, C.W., Isaias, I.U., Kusche-Tekin, B.B., Klein, D., Groh, J., O’Leary, A., Knorr, S., Higuchi, T., Koprich, J.B., Brotchie, J.M., et al. (2016). Tor1a+/- mice develop dystonia-like movements via a striatal dopaminergic dysregulation triggered by peripheral nerve injury. Acta Neuropathol Commun 4, 108.

140. Jinnah, H.A., Neychev, V., and Hess, E.J. (2017). The Anatomical Basis for Dystonia: The Motor Network Model. Tremor Other Hyperkinet Mov (N Y) 7, 506.

141. Balint, B., Mencacci, N.E., Valente, E.M., Pisani, A., Rothwell, J., Jankovic, J., Vidailhet, M., and Bhatia, K.P. (2018). Dystonia. Nat Rev Dis Primers 4, 25.

142. Egger, K., Mueller, J., Schocke, M., Brenneis, C., Rinnerthaler, M., Seppi, K., Trieb, T., Wenning, G.K., Hallett, M., and Poewe, W. (2007). Voxel based morphometry reveals specific gray matter changes in primary dystonia. Mov Disord 22, 1538–1542.

143. Bai, X., Vajkoczy, P., and Faust, K. (2021). Morphological Abnormalities in the Basal Ganglia of Dystonia Patients. Stereotact Funct Neurosurg, 1-12.

144. Vo, A., Nguyen, N., Fujita, K., Schindlbeck, K.A., Rommal, A., Bressman, S.B., Niethammer, M., and Eidelberg, D. (2023). Disordered network structure and function in dystonia: pathological connectivity vs. adaptive responses. Cereb Cortex 33, 6943–6958. 10.1093/cercor/bhad012.

145. Fremont, R., Tewari, A., Angueyra, C., and Khodakhah, K. (2017). A role for cerebellum in the hereditary dystonia DYT1. Elife 6.

146. Tewari, A., Fremont, R., and Khodakhah, K. (2017). It’s not just the basal ganglia: Cerebellum as a target for dystonia therapeutics. Movement disorders : official journal of the Movement Disorder Society 32, 1537–1545.

147. Liu, M.L., Zang, T., and Zhang, C.L. (2016). Direct Lineage Reprogramming Reveals Disease-Specific Phenotypes of Motor Neurons from Human ALS Patients. Cell reports 14, 115–128.

148. Pelossof, R., Fairchild, L., Huang, C.H., Widmer, C., Sreedharan, V.T., Sinha, N., Lai, D.Y., Guan, Y., Premsrirut, P.K., Tschaharganeh, D.F., et al. (2017). Prediction of potent shRNAs with a sequential classification algorithm. Nat Biotechnol 35, 350–353.

149. Dull, T., Zufferey, R., Kelly, M., Mandel, R.J., Nguyen, M., Trono, D., and Naldini, L. (1998). A third-generation lentivirus vector with a conditional packaging system. J Virol 72, 8463–8471.

150. Zufferey, R., Dull, T., Mandel, R.J., Bukovsky, A., Quiroz, D., Naldini, L., and Trono, D. (1998). Self-inactivating lentivirus vector for safe and efficient in vivo gene delivery. J Virol 72, 9873–9880.

151. Ding, B., and Kilpatrick, D.L. (2013). Lentiviral vector production, titration, and transduction of primary neurons. Methods Mol Biol 1018, 119–131.

152. Ding, B., Mirza, A.M., Ashley, J., and Budnik, V. (2017). Nuclear Export Through Nuclear Envelope Remodeling in Saccharomyces cerevisiae. bioRxiv *DOI:* 10.1101/224055.

153. Li, Y., Hassinger, L., Thomson, T., Ding, B., Ashley, J., Hassinger, W., and Budnik, V. (2016). Lamin Mutations Accelerate Aging via Defective Export of Mitochondrial mRNAs through Nuclear Envelope Budding. Curr Biol 26, 2052–2059.

154. Ding, B., and Kilpatrick, D.L. (2013). Chromatin immunoprecipitation assay of brain tissues using Percoll gradient-purified nuclei. Methods in molecular biology 1018, 199–209.

155. Ding, B., Cave, J.W., Dobner, P.R., Mullikin-Kilpatrick, D., Bartzokis, M., Zhu, H., Chow, C.W., Gronostajski, R.M., and Kilpatrick, D.L. (2016). Reciprocal autoregulation by NFI occupancy and ETV1 promotes the developmental expression of dendrite-synapse genes in cerebellar granule neurons. Mol Biol Cell 27, 1488–1499.

156. Ding, B., LeJeune, D., and Li, S. (2010). The C-terminal repeat domain of Spt5 plays an important role in suppression of Rad26-independent transcription coupled repair. J Biol Chem 285, 5317–5326.

157. Mi, H., Muruganujan, A., Casagrande, J.T., and Thomas, P.D. (2013). Large-scale gene function analysis with the PANTHER classification system. Nat Protoc 8, 1551–1566. 10.1038/nprot.2013.092.

158. Ding, B., Wang, W., Selvakumar, T., Xi, H.S., Zhu, H., Chow, C.W., Horton, J.D., Gronostajski, R.M., and Kilpatrick, D.L. (2013). Temporal regulation of nuclear factor one occupancy by calcineurin/NFAT governs a voltage-sensitive developmental switch in late maturing neurons. J Neurosci 33, 2860–2872.

159. Ding, B., Dobner, P.R., Mullikin-Kilpatrick, D., Wang, W., Zhu, H., Chow, C.W., Cave, J.W., Gronostajski, R.M., and Kilpatrick, D.L. (2018). BDNF activates an NFI-dependent neurodevelopmental timing program by sequestering NFATc4. Molecular biology of the cell 29, 975–987. 10.1091/mbc.E16-08-0595.

